# Motor primitives in space and time via targeted gain modulation in cortical networks

**DOI:** 10.1101/451054

**Authors:** Jake P. Stroud, Mason A. Porter, Guillaume Hennequin, Tim P. Vogels

## Abstract

Motor cortex (M1) exhibits a rich repertoire of activities to support the generation of complex movements. Although recent neuronal-network models capture many qualitative aspects of M1 dynamics, they can generate only a few distinct movements. Additionally, it is unclear how M1 efficiently controls movements over a wide range of shapes and speeds. We demonstrate that simple modulation of neuronal input–output gains in recurrent neuronal-network models with fixed architecture can dramatically reorganize neuronal activity and thus downstream muscle outputs. Consistent with the observation of diffuse neuromodulatory projections to M1, we show that a relatively small number of modulatory control units provide sufficient flexibility to adjust high-dimensional network activity using a simple reward-based learning rule. Furthermore, it is possible to assemble novel movements from previously learned primitives, and one can separately change movement speed while preserving movement shape. Our results provide a new perspective on the role of modulatory systems in controlling recurrent cortical activity.

## INTRODUCTION

Motor cortex is one of the final cortical outputs to downstream spinal motoneurons [1], and it is fundamental for controlling voluntary movements [2–4]. During movement execution, primary motor cortex (M1) exhibits complex, multiphasic firing-rate transients that return to baseline after movement completion [4]. Recent studies have provided some understanding of how these complex, single-neuron patterns of activity relate to intended movements [4–6]. It has been insightful to view motor cortex as a dynamical system in which preparatory activity sets the initial condition for the system, whose subsequent dynamics drive the desired muscle activity [7, 8]. From this perspective, the complex firing-rate dynamics provide a flexible basis set for the generation of movements [9].

Several recurrent neuronal-network models have been developed to capture M1 activity during movement execution [10, 11]. These models rely on strong recurrent connectivity that is optimized for the neuronal dynamics to be qualitatively similar to M1 activity during movement execution. However, these models cannot explain how new movements can be constructed or how their static architecture allows variations in both output trajectories and speed.

A possible mechanism for effectively switching neuronal activity, and consequently downstream muscle activity to generate different movements (see Fig. 1a), is to adjust the intrinsic gain — that is, the input–output sensitivity — of each neuron so that they engage more (or less) actively in the recurrent neuronal dynamics [12–18]. Indeed, neuromodulation in M1 can cause such changes in neuronal responsiveness [19, 20], and gain modulation of both neurons in M1 [13] and spinal motoneurons [21, 22] has been linked experimentally to skill acquisition and optimization of muscular control.

**Figure 1:**
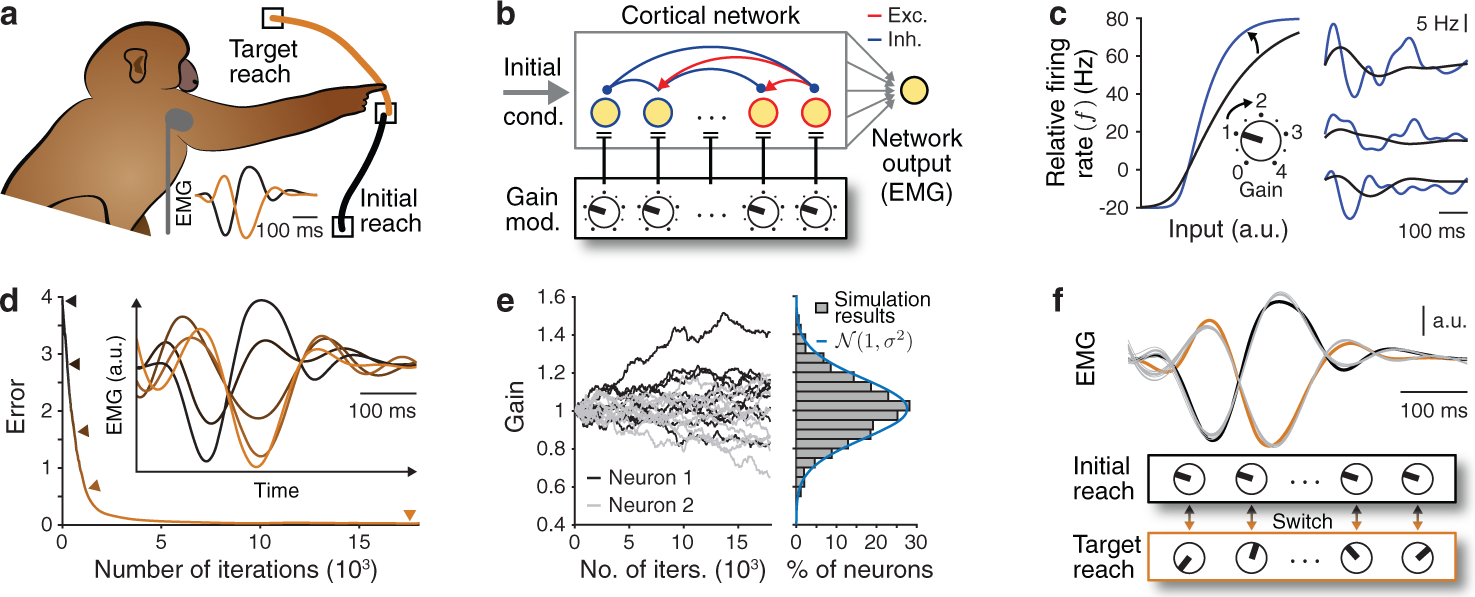
Controlling network activity through neuron-specific gain modulation. (**a**) Example of a reaching task, with illustrative electromyograms (EMGs) of muscle activity for two reaches (in orange and black). (**b**) Schematic of our model (see the text and Methods Section 1.10). (**c**) (Left) Changing the slope of the input–output gain function uniformly for all neurons from (black) 1 to (blue) 2 has pronounced effects on (right) neuronal firing rates. We show results for three example neurons. (**d**) The mean error in network output decreases during training with neuron-specific modulation. In the inset, we show five snapshots of network output (indicated by arrowheads) as learning progresses. (**e**) (Left) Neuronal gain changes during training for 2 example neurons (grey and black) and 10 training sessions. (Right) Histogram of gain values after training. The blue curve is a Gaussian fit with a standard deviation of *σ* ≈ 0.157. (**f**) Network outputs (grey curves) with all gains set to 1 and the new learned gain pattern for 10 noisy initial conditions compared to both targets (black and orange). (We use a 200-neuron network for all simulations in this figure.)

In this paper, we study the effects of gain modulation in recurrent neuronal-network models of motor cortex. We show that individually modulating the gain of neurons in such models allows learning of a variety of target outputs on behaviourally relevant time scales through reward-based training. Motivated by diffuse neuromodulatory innervation of M1 [19, 23, 24], we find that coarse-grained control of neuronal gains achieves a similar performance to neuron-specific modulation. We demonstrate that we can combine previously learned modulatory patterns to accurately generate new desired movements. Therefore, gain patterns can act as motor primitives for quickly constructing novel movements [25, 26]. Finally, we show how to control the speed of an intended movement through gain modulation. We find that it is possible to learn gain patterns that affect either only the shape or only the speed of a movement, thus enabling efficient and independent movement control in space and time.

## RESULTS

### Modelling gain modulation in recurrent neuronal networks

To understand how a cortical network can efficiently generate a large variety of outputs, we begin with an existing cortical circuit model [11]. We use recurrent networks, with *N* = 2*M* neurons (where *M* are excitatory and *M* are inhibitory), for which the neuronal activity vector *x*(*t*) = (*x*_1_(*t*), …, *x_N_*(*t*))^┬^ evolves according to

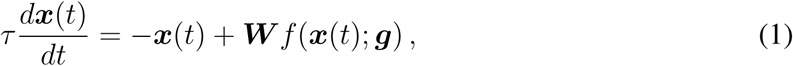

where the single-neuron time constant is *τ* = 200 ms, and (unless we state otherwise) we generate the synaptic weight matrix *W* in line with [11] (i.e., we use ‘stability-optimized circuits’). These networks consist of a set of sparse, strong excitatory weights that are balanced by fine-tuned inhibition (see Methods).

The gain function *f*, which governs the transformation of neuronal activity *x* into firing rates relative to a baseline rate *r*_0_, is

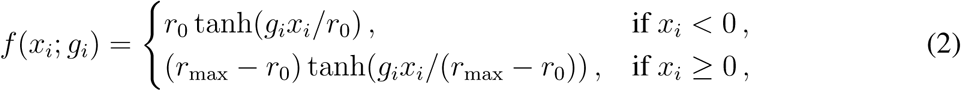

where *g_i_* is the slope of the function *f* at the baseline rate *r*_0_ and thus controls the input–output sensitivity of neuron *i* at spontaneous activity levels [27]. In Eqn. (1),*f* (*x*;*g*) denotes the element-wise application of the scalar function *f* to the neuronal activity vector *x*. Unless we state otherwise, we use a baseline rate of *r*_0_ = 20 Hz and a maximum firing rate of *r*_max_ = 100 Hz, consistent with experimental observations [4, 28]. With this setup,*f* (*x*;*g*) describes the neuronal firing rates relative to the baseline steady-state *r*_0_. We also note that identical dynamics can result from using a strictly positive gain function, combined with a tonic (i.e., static) external input (see Methods Section 1.2).

For appropriate initial conditions *x*(*t* = 0) = *x*_0_ (see Methods Section 1.1), the neuronal dynamics given by Eqn. (1) exhibit naturalistic activity transients that resemble M1 recordings [4, 11], and the population activity is rich enough to enable the generation of complex movements through linear readouts [11]. We emulate neuromodulation in this model by directly controlling the input–output gain *g_i_* of each neuron (see Figs. 1b,c).

### Neuron-specific gain modulation

We find that increasing the gain of all neurons uniformly (i.e., *g_i_* = *g* in Eqn. (2)) increases both the frequency and amplitude of neuronal firing rates (see Fig. 1c). One can understand these effects of uniform modulation by linearizing Eqn. (1) around *x* = 0, yielding the linear ordinary differential equation 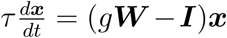 (where *I* is the identity matrix), and studying changes in the spectrum of the matrix *g**W** − **I***; see our Supplementary Maths Note.

To allow more precise control of network activity than through uniform modulation, we can independently adjust the gain of each neuron in what we call *neuron-specific modulation*. We obtain gain patterns that lead to the generation of target output activity using a reward-based node perturbation learning rule (see Methods Section 1.7). Our rule, which acts on the modulatory pathway of our model but is similar to proposed synaptic plasticity rules for reward-based learning [29–32], uses a global scalar signal of recent performance to iteratively adjust each neuron’s gain while the initial condition *x*_0_ and the network architecture remain fixed.

Starting with a network and readout weights that produces an initial movement with all gains set to 1 (see the black curve in Fig. 1d), our learning rule yields a gain pattern that leads to the successful generation of a novel target movement after a few thousand training iterations (see Fig. 1d and Methods Section 1.10). Errors between the actual and desired outputs tend to decrease monotonically and eventually become negligible. Independent training sessions with the same target movement produce nonidentical but positively correlated gain patterns (see Fig. 1e and Supplementary Fig. 1c). Counterintuitively, the neuronal firing rates change only slightly, even though the network output is altered substantially (see Supplementary Figs. 1b). Once the target is learned, the same initial condition can produce either of two distinct network outputs, depending on the applied gain pattern (see Fig. 1f). The outputs are also similarly robust with respect to noisy initial conditions for each gain pattern (see Supplementary Fig. 1d).

We also compare the learning performance of gain modulation with alternative learning mech-anisms. We train either the neuronal gains, the initial condition *x*_0_ of the neuronal activity, a rank-1 perturbation of the synaptic weight matrix, or the full synaptic weight matrix using back-propagation (see Methods Section 1.10). We find empirically for this task that training through gain modulation yields a similar learning performance as training the initial condition or the full synaptic weight matrix and that training through gain modulation performs significantly better than learning a rank-1 perturbation of the synaptic weight matrix (see Supplementary Fig. 1f).

### Gain modulation in different models

We next examine whether learning through gain modulation is possible in alternative, commonly used variants of our model [10, 11]. Motor circuits that drive movements also engage in periods of movement preparation [5, 7, 33], suggesting a role for gain modulation in shaping circuit dynamics both during movement planning and during movement execution. We find that learning is also possible in a model in which we include gain modulation during movement planning. We simulate the preparatory period using a ramping input to the system [11] (see Methods Section 1.10), such that gain modulation now directly affects the neuronal activity at movement onset. We find that learning performance (i.e., error reduction) for the task that we showed in Fig. 1d is slightly poorer if we do employ a ramping input than if we do not. (Compare the red and blue curves in Fig. 2a.)

**Figure 2:**
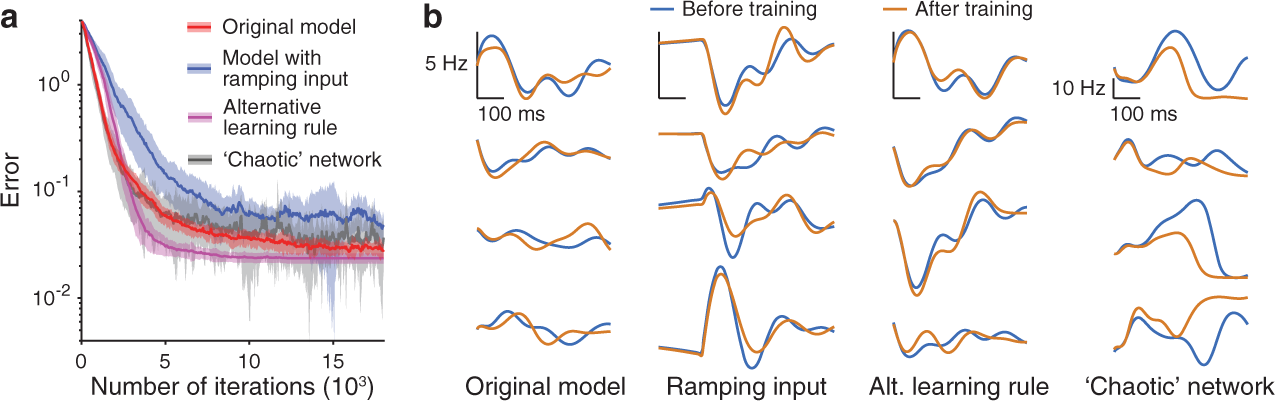
Learning through gain modulation in different models. (**a**) Mean error over 10 independent training sessions for our original model from Fig. 1d (red); the model with a biologically motivated ramping input (blue); the model when using the alternative learning rule Eqn. (10) in which learning automatically stops at a sufficiently small error (purple); and when using a ‘chaotic’ recurrent network model (grey) (see Methods Section 1.10). Shading indicates one standard deviation. (**b**) The firing rates of 4 example neurons before (i.e., with all gains set to 1) and after training the neuronal gains in (left) our original model, (centre left) our model with a ramping input, (centre right) our model with the alternative learning rule, and (right) the model when using a ‘chaotic’ network. (We use 200-neuron networks for all simulations in this figure.)

This occurs because gain modulation during the preparatory phase changes the neuronal activity at movement onset, allowing it to leave the null space of the readout weights (which are fixed) and thus elicit premature muscle activity at movement onset.

We also construct a ‘chaotic’ variant of our model [34] (see Methods Sections 1.3 and 1.10) for the same task and train only the neuronal gains. We achieve similar learning performance compared to our original model from Fig. 1d (compare the red and grey curves in Fig. 2a), even though the neuronal dynamics are very different (compare the far left and far right panels in Fig. 2b). Finally, we also use an alternative learning rule to train the neuronal gains (see Eqns. (10) and (11)); in this rule, learning slows down as the decrease in error slows down (see Methods Section 1.8). We find that the error decreases at a faster rate than that in the original learning rule (see the purple curve in Fig. 2a.) This may occur because the standard deviation of the noise perturbation term in the alternative learning rule becomes smaller over training iterations as the error decreases.

Notably, in all of these scenarios, changes in neuronal responsiveness alone — for example, via inputs from neuromodulatory afferents — can cause dramatic changes in network outputs, thereby providing an efficient mechanism for rapid switching between movements, without requiring any changes in either synaptic architecture or the initial condition *x*_0_.

### Coarse, group-based gain modulation

Individually modulating the gain of every neuron in motor cortex is likely unrealistic. In line with the existence of diffuse (i.e., not neuron-specific) neuromodulatory projections to M1 [19, 23, 24], we cluster neurons into groups so that we identically modulate units within a group. (See Fig. 3a and Methods Sections 1.9 and 1.10.) We find that such coarse-grained modulation gives similar performance to neuron-specific control for as few as 20 randomly formed groups (see Methods Section 1.9) using our 200-neuron network model from Fig. 1. (See Fig. 3b and Supplementary Fig. 2a.) For a specified number of groups, one can improve performance if, instead of grouping neurons randomly as above, we use a specialized clustering for each movement that is based on previous training sessions (see Fig. 3b, Supplementary Fig. 2a, and Methods Section 1.9). Importantly, there exist specialized groupings that perform similarly across multiple different movements. (See Fig. 3c and Supplementary Figs. 2b,c.) Such specialized groupings acquired from learning one set of movements also perform well on novel movements (see Supplementary Fig. 2d).

**Figure 3:**
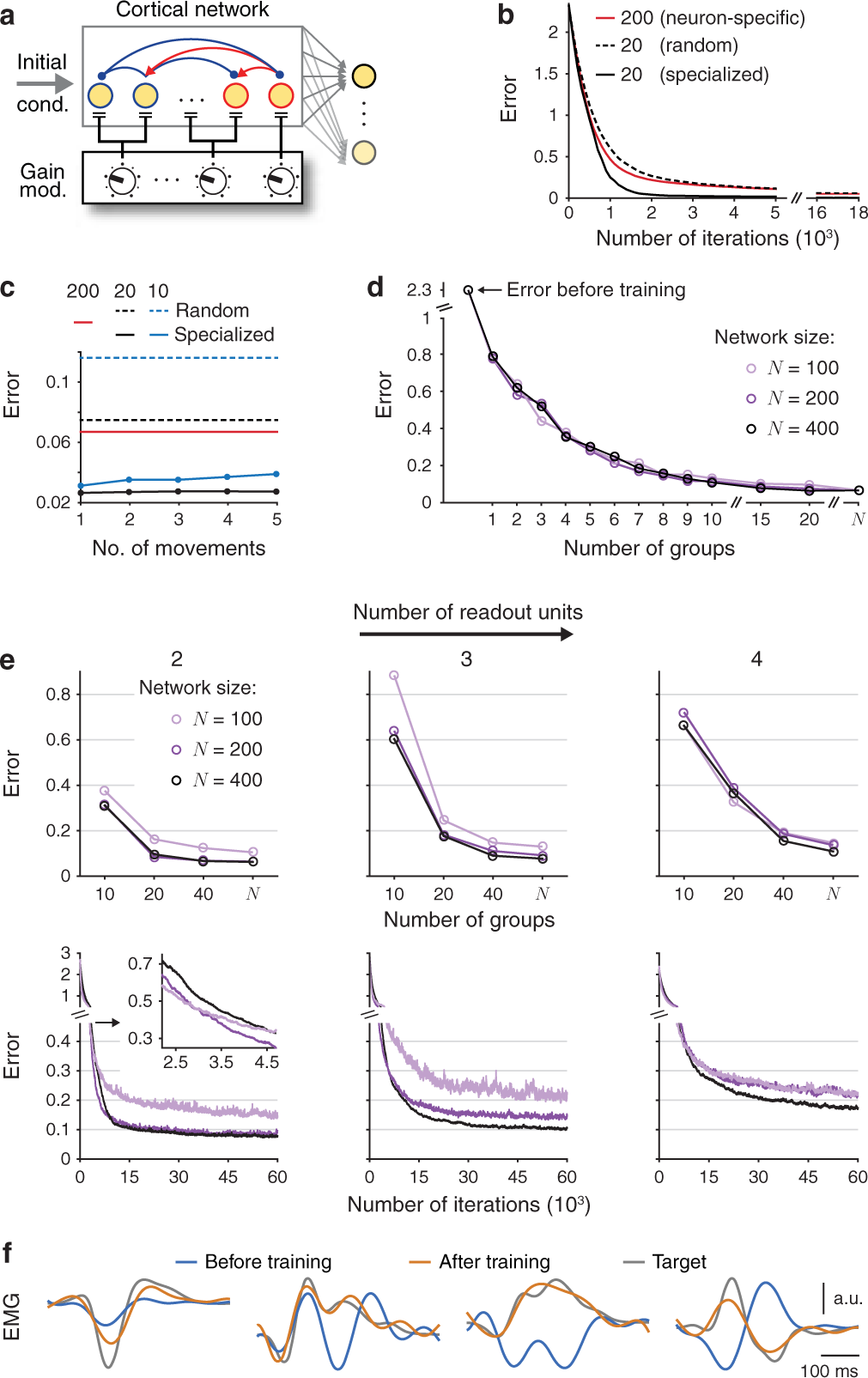
Controlling network activity through coarse, group-based gain modulation. (**a**) We identically modulate neurons within each group (see Methods Section 1.9). Target outputs can involve multiple readout units. (**b**) Mean error during training for 20 random, 20 specialized, and 200 (i.e., neuron-specific) groups. (See Methods Section 1.10 for precise specifications of these examples.) (**c**) Mean minimum errors after training using specialized groups. We use the same groupings for different numbers of movements. In panels b–c we use a 200-neuron network. (**d**) Mean minimum errors for different numbers of random groups with networks of 100, 200, and 400 neurons. (The *N* on the horizontal axis indicates neuron-specific modulation.) In panels (b)–(d), we use a single readout unit. (**e**) (Top) Mean minimum error as a function of the number of random groups when learning each of (left) 2, (centre) 3, and (right) 4 readouts for the same networks as in panel (d). (Bottom) The corresponding mean errors during training for the case of 40 groups. The inset is a magnification of the initial training period for the case of 2 readout units. (**f**) Outputs producing the median error for the case of 4 readout units using 40 groups in the 400-neuron network.

Notably, even with random groupings, network size hardly affects learning performance for a single readout (see Fig. 3d). Performance depends much more on the number of groups than on the number of neurons per group. When the task involves two or more readout units, larger networks do learn better, and achieving a good performance necessitates using a larger number of independently modulated groups (see Fig. 3e). Finally, smaller networks typically learn faster (see the bottom panel of Fig. 3e), but they ultimately exhibit poorer performance, demonstrating that there is a trade-off between network size, number of groups, and task complexity (i.e., the number of readout units).

### Gain patterns can provide motor primitives for novel movements

In principle, it is possible to independently learn numerous gain patterns, supporting the possibility of a repertoire (which we call a ‘library’) of modulation states that a network can use, in combination, to produce a large variety of outputs. Generating new movements is much more efficient if it is possible to ‘intuit’ new gain patterns as combinations of previously acquired primitives [15, 26]. To test if this is possible in our model, we first approximate a novel target movement as a convex combination of existing movements. (We call this a ‘fit’ in Fig. 4; see Methods Section 1.10.) We then use the same combination of the associated library of gain patterns to construct a new gain pattern (see Fig. 4a). Interestingly, the resulting network output closely resembles the target movement (see Fig. 4b). This may seem unintuitive, but one can understand this result mathematically by calculating power-series expansions of the solution of the linearized neuronal dynamics (see our Supplementary Maths Note).

**Figure 4:**
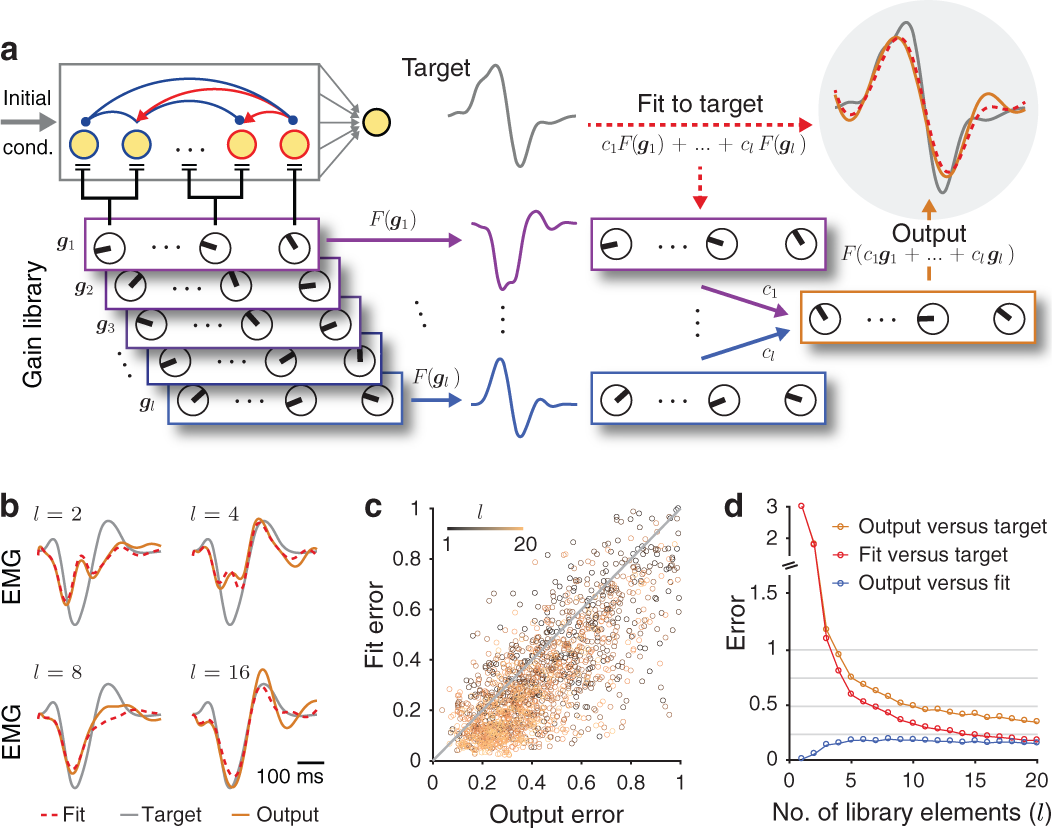
Gain patterns can provide motor primitives for novel movements. (**a**) Schematic of a learned library of gain patterns (*g*_1_, …, *g_l_*, which we colour from purple to blue) and a combination *c*_1_*F* (*g*_1_) + … + *c_l_F*(*g_l_*) of their outputs (which we denote by *F*) that we fit (red dashed curve) to a novel target (grey curve). (Upper right) The output *F*(*c*_1_*g*_1_ + … + *c_l_g_l_*) (which we show in orange) of the same combination of corresponding gain patterns also closely resembles the target. We use a 400-neuron network with 40 random modulatory groups (see Methods Section 1.10). (**b**) Example target, fit, and output (grey, red dashed, and orange curves, respectively) producing the 50th-smallest output error over 100 randomly generated combinations (see Methods Section 1.10) of *l* library elements using *l* = 2, *l* = 4, *l* = 8, and *l* = 16. (**c**) Fit error versus the output error for 100 randomly generated combinations of *l* library elements for *l* = 1, …, 20. We show the identity line in grey. Each point represents the 50th-smallest error between the output and the fit across 100 novel target movements. (**d**) Median errors of the 100 randomly generated combinations of *l* library elements versus the number of library elements.

Finally, increasing the number of elements in the movement library reduces the error between a target movement and its fit, which is also reflected in a progressively better match between the target and the network output (see Figs. 4b–d and Supplementary Fig. 3). Although the idea of using motor primitives to facilitate rapid acquisition of new movements is well established [25, 26], our approach proposes the first (to our knowledge) circuit-level mechanism for achieving this objective. In addition to neuromodulatory systems [19, 20, 22], the cerebellum is a natural candidate structure to coordinate such motor primitives [25], as it is known to project to M1 and to play a critical role in error-based motor learning [25, 35].

### Nonlinear behaviour

We initially choose the baseline firing rate (*r*_0_ = 20 Hz in Eqn. (2)) to be consistent with experimentally measured firing rates in motor cortex [4, 28, 36]. Most of the time, neurons operate within the linear part of their nonlinear gain function *f* (i.e., the neuronal dynamics are similar to the case of using the linear gain function *f* (*x_i_; g_i_*) = *g_i_x_i_* (see Figs. 5a,c)). To test if our results hold for scenarios with more strongly nonlinear dynamics, we reduce the baseline firing rate to *r*_0_ = 5 Hz. This increases the neuronal activity near the lower-saturation regime (i.e., towards the left part of the curve in the left panel of Fig. 1c) of the gain function (see Figs. 5b,c). As expected from the larger range of possible network outputs (and improved learning performance) in nonlinear recurrent neuronal networks than in linear ones [31, 32, 34], we observe better learning performance for *r*_0_ = 5 Hz than for *r*_0_ = 20 Hz (compare the black and blue curves in Fig. 5d), and we obtain a very similar distribution of gain values after training (see Fig. 5e).

**Figure 5:**
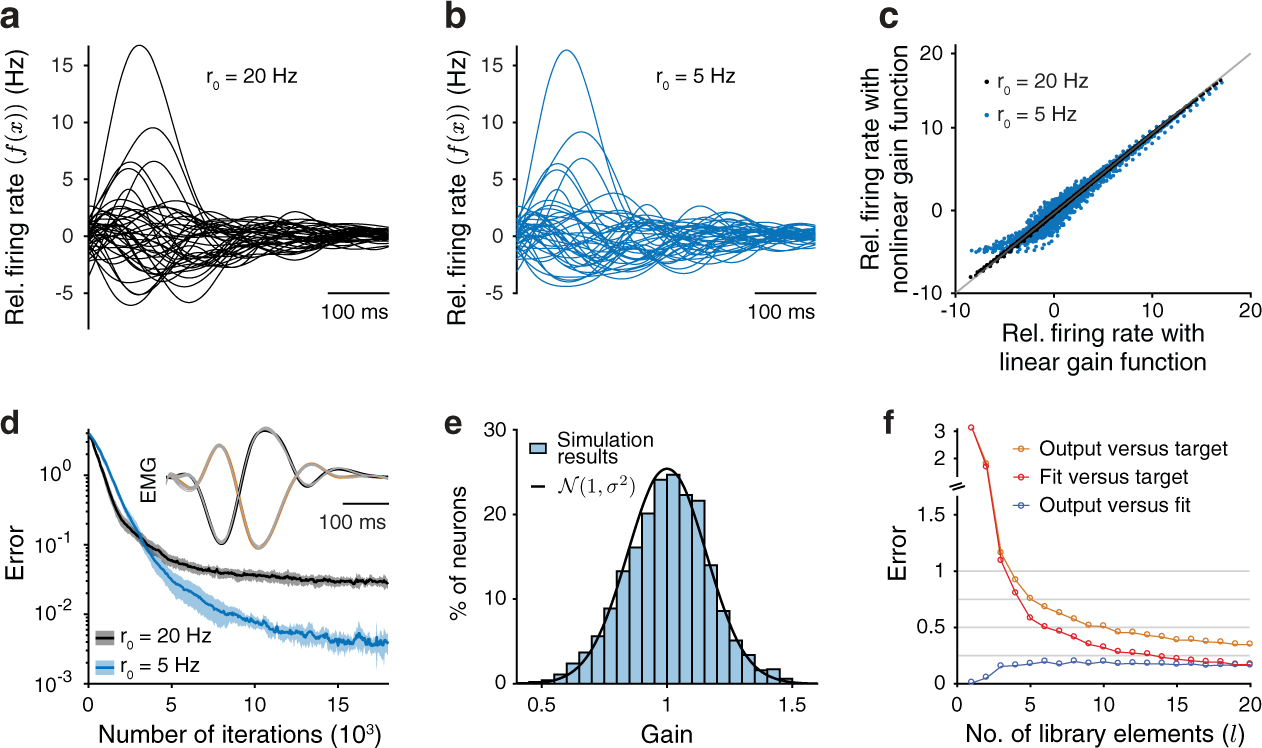
Examining effects of more strongly nonlinear neuronal dynamics by using a baseline rate of *r*_0_ = 5 Hz. (**a**) Relative firing rate of 20 excitatory and 20 inhibitory neurons in a 200-neuron network with *r*_0_ = 20 Hz in Eqn. (2). (**b**) Relative firing rate of the same neurons as those in panel (a), but with *r*_0_ = 5 Hz. (**c**) The dotted curves show the relative firing rates of all neurons over time when using the nonlinear gain function *f* (see Eqn. (2)) with (black) *r*_0_ = 20 Hz and (blue) *r*_0_ = 5 Hz versus the relative firing rates that result from using the linear gain function *f*(*x_i_; g_i_*) = *g_i_x_i_* in Eqn. (1). We set each neuronal gain to 1, and we plot the identity line in grey. (**d**) Mean error over 10 independent training sessions with *r*_0_ = 20 Hz (black) and with *r*_0_ = 5 Hz (blue) for the task in Fig. 1d (see Methods Section 1.10). Shading indicates one standard deviation. In the inset, we show network outputs with all gains set to 1 and the new learned gain pattern with *r*_0_ = 5 Hz for 10 noisy initial conditions (grey curves). We show the two targets in black and orange (see Methods Section 1.10). (e) Histogram of gain values after training with *r*_0_ = 5 Hz. The black curve is a Gaussian distribution with mean 1 and standard deviation *σ* ≈ 0.157 (i.e., the distribution that we obtained with *r*_0_ = 20 Hz in Fig. 1e). (f) Gain patterns as motor primitives with *r*_0_ = 5 Hz. We generate these results in the same manner as our results in Fig. 4d, except that now we use *r*_0_ = 5 Hz. We obtain qualitatively similar results to our observations for the baseline rate *r*_0_ = 20 Hz.

Importantly, it is still possible to learn new movements by using combinations of existing gain patterns. As before, performance is limited by the accuracy with which one can construct target movements as linear combinations of existing primitives. (See the correlations between network output errors and fit errors in Supplementary Fig. 4b.) Moreover, errors in network output decrease on average with increasing numbers of gain patterns in the movement library (see the orange curve in Fig. 5f), and the difference between the network output and corresponding fit remains small for all tested numbers of library elements (see the blue curve in Fig. 5f). However, reducing *r*_0_to sufficiently small values (that are below 5 Hz) does eventually lead to a deterioration in the relationship between gain patterns and their corresponding outputs.

### Gain modulation can control movement speed

Thus far, we have demonstrated that simple (even coarse, group-based) gain modulation enables control of network outputs of the same, fixed duration. To control movements of different durations, motor networks must be able to slow down or speed up muscle outputs (i.e., change the duration of movements without affecting their shape). In line with recent experimental results [37, 38], we investigate if changing neuronal gains allows control of the speed of an intended movement. (See Fig. 6a and Methods Section 1.10.) We begin with a network of 400 neurons (with 40 random modulatory groups) that generates muscle activity that lasts approximately 0.5 s (as in Figs. 3 and 4). We find that our learning rule can successfully train a network to generate a slower variant that lasts 5 times longer than the original movement (see Fig. 6b, Supplementary Fig. 5a, and Methods Section 1.10).

**Figure 6:**
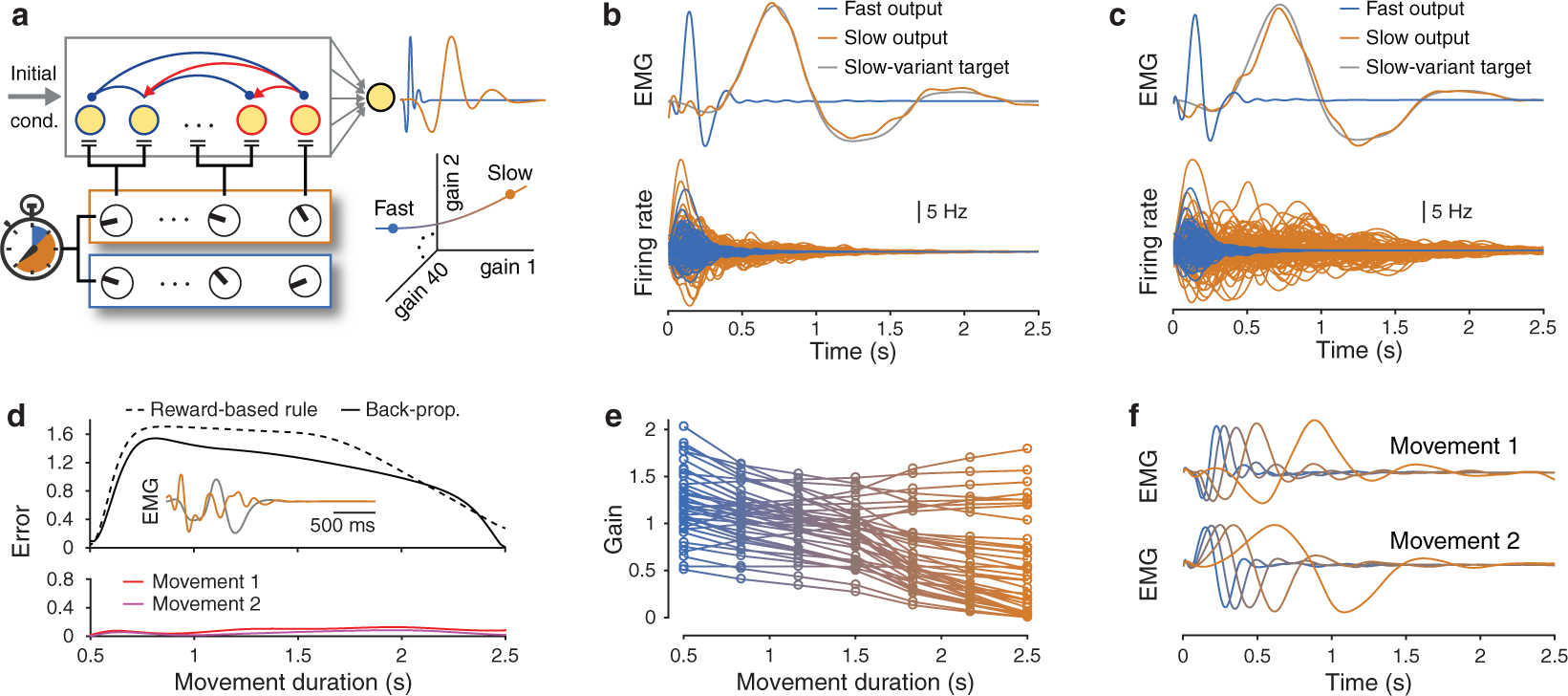
Gain modulation can control movement speed. (**a**) Schematic of gain patterns for fast (0.5 s) and slow (2.5 s) movement variants. (Here and throughout the figure, we show the former in blue and the latter in orange.) We train a 400-neuron network using 40 random modulatory groups for all simulations (see Methods Section 1.10). (**b**) (Top) We train a network to extend its output from a fast to a slow-movement variant using our reward-based learning rule. (Bottom) Example firing rates of 50 excitatory and 50 inhibitory neurons for both fast and slow speed variants. (**c**) The same as panel (b), but now we use a back-propagation algorithm to train the neuronal gains (see Methods Section 1.10). (**d**) (Top) Interpolation between fast and slow gain patterns does not reliably generate target outputs of intermediate speeds when trained only at the fast and slow speeds. We show an example output (orange) that lasts a duration of 1.5 s and the associated target (grey). (Bottom) Linear interpolation between the fast and slow gain patterns successfully generates target outputs when trained at 5 intermediate speeds. We train 1 set of gain patterns (see panel (e)) on two target outputs associated with 2 different initial conditions (see Methods Section 1.10). (We plot these results with the same axis scale as in the top panel.) (**e**) The 7 optimized gain patterns for all 40 modulatory groups when training at 7 evenly-spaced speeds. (**f**) Both outputs for 5 evenly-spaced speeds between the fast and slow gain patterns from panel (e).

In contrast to simply changing the single-neuron time constant *τ* — which uniformly scales the duration, but does not affect the shape of each neuron’s activity — modifying neuronal gains to generate ‘fast’ and ‘slow’ output variants leads to changes in both the shape and duration of neuronal activities, in line with recent experimental findings [37]. Changing neuronal gains thus enables interactions between the shape and duration of outputs without requiring retraining of the synaptic weight matrix to scale the duration of neuronal activities [39].

The learned slow variants are more sensitive to noisy initial conditions than the fast variants, but we can find more robust solutions by using a regularized back-propagation algorithm to train both the neuronal gains and the readout weights (see Methods Section 1.10). Following training, the slow variants are learned successfully (see Fig. 6c) and are less sensitive to the same noisy initial conditions (see Supplementary Fig. 5g). The neuronal dynamics oscillate transiently, with a substantially lower frequency than either the fast variants or the slow variants trained by our reward-based learning rule. (Compare the bottom panels of Fig. 6c and Fig. 6b.) We also find a single gain pattern that, rather than slowing down only one movement, can slow down up to approximately five distinct movements, which result from five orthogonal initial conditions, by a factor of 5 (see Supplementary Figs. 5h–j). Consequently, one can extend the temporal scale of transient neuronal activity several-fold through specific changes in neuronal gains.

### Smoothly controlling the speed of movements

Following training on a slow and a fast variant of the same movement (see the previous section), we find that naively interpolating between the two gain patterns does not yield the same movement at intermediate speeds (see the top panel of Fig. 6d), consistent with human subjects being unable to consistently apply learned movements at novel speeds [39, 40]. Therefore, even when we consider ‘fast’ and ‘slow’ variants of the same movement, both our learning rule and the back-propagation training do not learn to ‘slow down’ the movement; instead, they learn two seemingly unrelated gain patterns. However, it is possible to modify our back-propagation training procedure by including additional constraints on the fast and slow gain patterns (see Methods Section 1.10) so that interpolating between the two gain patterns produces progressively faster or slower outputs. We successfully train the network to generate two movements (associated with two different initial conditions) at 7 different speeds with durations that range from 0.5 s to 2.5 s. (See Fig. 6e, Supplementary Fig. 6, and Methods Section 1.10.) Linear interpolation between the fast and slow gain patterns (see Supplementary Fig. 6b) now generates smooth speed control of both movements at any intermediate speed (see the bottom panel of Fig. 6d as well as Fig. 6f). In other words, to control movement speed, we learn a ‘manifold’ [41] in neuronal gain space that interpolates between the fast and the slow gain patterns. (See the bottom right of Fig. 6a.)

### Joint control of movement shape and speed

Thus far, we have shown that gain modulation can affect either the shape or the speed of a movement. Flexible and independent control of both the shape and speed of a movement (i.e., joint control) necessitates separate representations of space and time in the gain patterns. A relatively simple possibility is to find a single universal manifold in neuronal gain space (see the previous subsection) for speed control (we call this the ‘speed manifold’) and combine it with gain patterns that are associated with different movement shapes. Biologically, this may be achievable using separate modulatory systems. We achieve such separation by simultaneously training one speed manifold and 10 gain patterns for 10 different movement shapes such that movements are encoded by the product of shape-specific and speed-specific gain patterns. (See Fig. 7a and Methods Section 1.10.) Following training, we can generate each of the 10 movements at the 7 trained speeds by multiplying a speed-specific gain pattern (see Fig. 7b) with the desired shape-specific gain pattern. Importantly, we can also accurately generate each of the 10 different movements at any intermediate speed by simply linearly interpolating between the fast and slow gain patterns (see Figs. 7c,d). We thereby obtain separate families of gain patterns for movement shape and speed that independently control movements in space and time.

**Figure 7:**
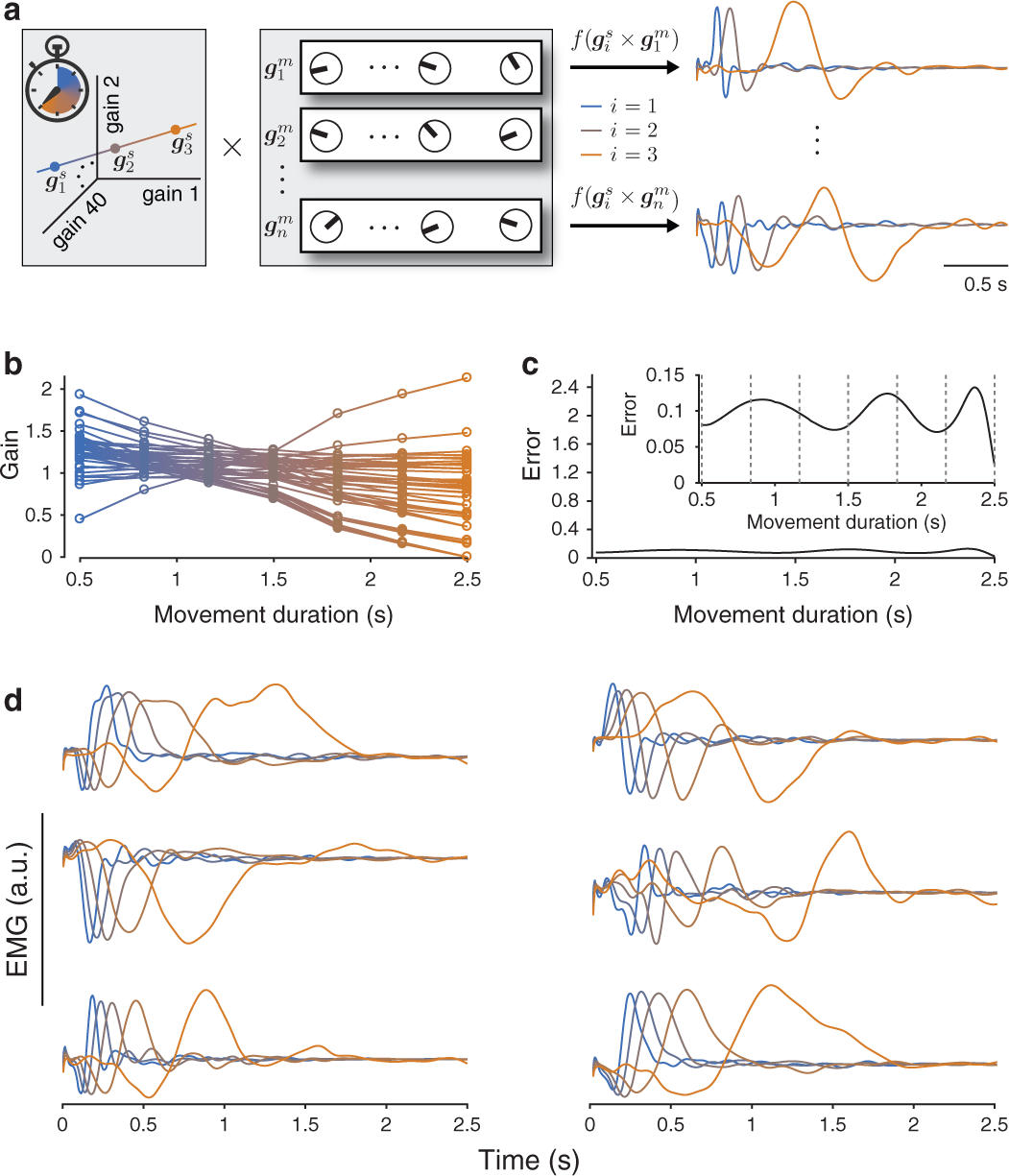
Joint control of movement shape and speed through gain modulation. (**a**) One can jointly learn the gain patterns 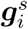 for (left box) movement speed and 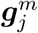 for (right box) movement shape so that the product of two such gain patterns produces a desired movement at a desired speed. In the rightmost panel, we show example outputs for two movement shapes at 3 interpolated speeds between the fast and slow gain patterns. (See the main text and Methods Section 1.10.) (b) We show the 7 optimized gain patterns for controlling movement speed (i.e., 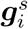 for *i* ∈ {1, …,7} from panel (a)) for the 40 modulatory groups when training on 10 different movement shapes. (c) We plot the mean error over all 10 movements when linearly interpolating between the fast and slow gain patterns for controlling movement speed from panel (b). We use the same vertical axis scale as in Fig. 6d. In the inset, we plot the same data using a different vertical axis scale. The vertical dashed lines identify the 7 movement durations that we use for training. (d) Outputs at 5 interpolated speeds between the fast and slow gain patterns for 6 of the 10 movements. (For each simulation, we train a 400-neuron network using 40 random modulatory groups (see Methods Section 1.10).)

### Learning gain-pattern primitives to control movement shape and speed

To construct new movement shapes with arbitrary durations, we examine the possibility of using both the speed manifold and the 10 trained shape-specific gain patterns that we obtained previously (see Fig. 7) as a library of spatiotemporal motor primitives. We test this library using 100 novel target movement shapes (see Fig. 4). For each target, we learn the coefficients for linearly combining the 10 shape-specific gain pattern primitives to obtain each new movement at both the fast and slow speeds while keeping the speed manifold fixed. (See Fig. 8a and Methods Section 1.10.)

**Figure 8:**
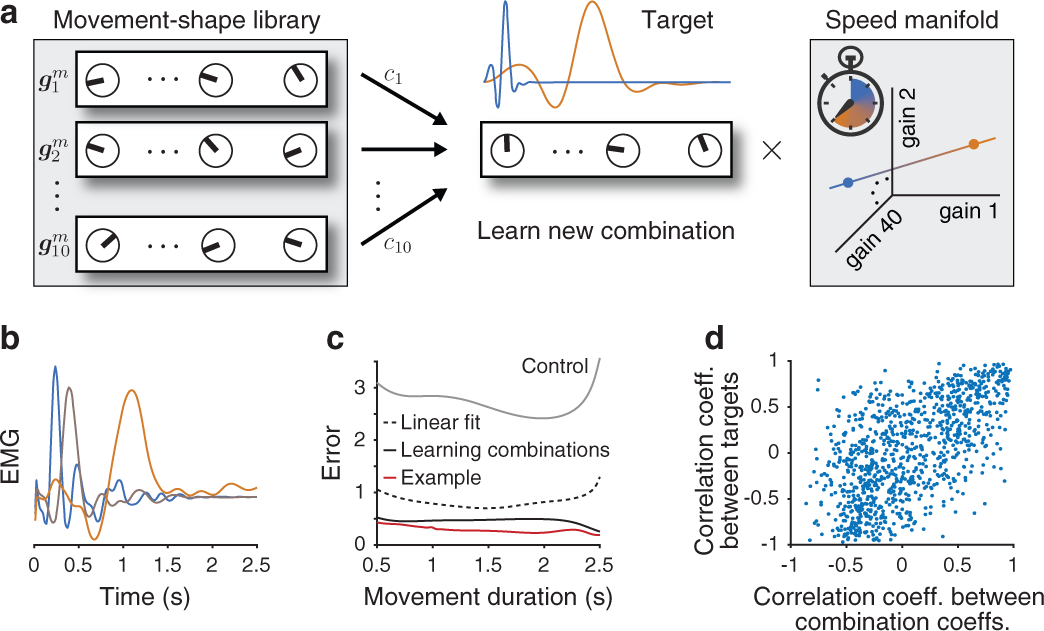
Learning gain-pattern primitives to control movement shape and speed. (**a**) We are able to learn to combine (left) previously acquired gain patterns for movement shapes to generate (centre) a new target movement at both fast and slow speeds simultaneously using (right) a fixed manifold in neuronal gain space for controlling movement speed (see Methods Section 1.10). (**b**) We plot the output, at 3 different speeds, that produces the 50th-smallest error (across all 100 target movements) between the output and the target when summing errors at both fast and slow speeds. (**c**) Mean network output error across all 100 target movements for all durations when learning to combine gain patterns (black solid curve). We plot the error for the output from panel (b) in red. As a control, we plot the mean error over all target movements when dissociating the learned gain patterns from their target movement by permuting (uniformly at random) the target movements (see the grey curve). We also plot the mean error over all target movements when combining gain patterns using a least-squares fit of the 10 learned movement shapes to the target (black dashed curve) (see Methods Section 1.10). (To generate outputs of a specific duration, we linearly interpolate between the fast and slow gain patterns.) (**d**) We plot the Pearson correlation coefficient between each pair of target movements versus the Pearson correlation coefficient between the corresponding pair of learned combination coefficients *c*_1_, …, *c*_10_. (For each simulation, we train a 400-neuron network using 40 random modulatory groups (see Methods Section 1.10).)

We find that it is possible to accurately generate the new movements at fast and slow speeds using the above spatiotemporal library of gain patterns (see Supplementary Fig. 7). Importantly, we are able to produce the new movements with similar accuracies as those at the fast and slow speeds at any intermediate speed by linearly interpolating between the fast and slow gain patterns using the unaltered speed manifold. (See Fig. 8b and the black and red curves in Fig 8c.) The mean error of approximately 0.5 across all movement durations is similar to the error that we obtained previously from a movement library that consists of 10 gain patterns (see Fig. 4d). We can substantially outperform both the (uniformly-at-random) permuted gain patterns from their associated targets (see Methods Section 1.10) and using least-squares fitting (which we used previously) to combine gain patterns. (See the grey and black dashed curves in Fig. 8c.)

Consistent with the idea of rapidly generating movements using motor primitives, we generate correlated target shapes by using correlated combinations of gain patterns (see Fig. 8d). Therefore, one can use previously learned gain patterns for controlling movement shapes to generate new movements while maintaining independent control of movement speed.

## DISCUSSION

The movement-specific population activity that has been observed in monkey primary motor cortex [4], can arise through several possible mechanisms. Distinct neuronal activity can emerge from a fixed population-level dynamical system with different movement-specific preparatory states [7]. Alternatively, one can change the underlying dynamical system through modification of the effective connectivity [42] even when a preparatory state is the same across movements. Such changes in effective connectivity can arise either through a feedback loop (e.g., a low-rank addition to the synaptic weight matrix [34]) or through patterns of movement-specific gains, as we explored in this paper. We found that movement-specific gain patterns provide a similar performance to training a different initial condition for each desired output (with a fixed duration) and that both of these approaches outperform a rank-1 perturbation of the synaptic weight matrix (see Supplementary Fig. 1f). Gain modulation thus provides a complementary method of controlling neuronal dynamics for flexible and independent manipulation of output shape. Additionally, gain modulation provides a compelling mechanism for extending the duration of activity transients without needing to carefully construct movement-specific network architectures [39].

Gain modulation can occur primarily via neuromodulators [20, 22], but it can also arise from a tonic (i.e., static) input that shifts each neuron’s resting activity within the dynamic range of its input–output function (for example, through inputs from the cerebellum) [14]. Although this is an effective way of mimicking gain changes in recurrent network models with strongly nonlinear single-neuron dynamics [37, 43], we were unable to produce desired target outputs by training a tonic input. It is worth noting that a tonic input also modifies baseline neuronal activity, thereby altering the output muscle activities away from rest.

In our model, in which the recurrent architecture remains fixed, synaptic modifications can take place upstream of the motor circuit (e.g., in the input synapses to the presumed neuromodulatory neurons [44]). Additionally, changes in neuronal gains can work in concert with plasticity in cortical circuits, thereby allowing changes in the modulatory state of a network to be transferred into circuit connectivity [45], consistent with known interactions between neuromodulation and plasticity [44]. Consequently, understanding the neural basis of motor learning may necessitate recording from a potentially broader set of brain areas than those circuits whose activity correlates directly with movement dynamics.

Our results build on a growing literature of taking a dynamical-systems approach to studying temporally structured cortical activity. This perspective has been effective for investigations of several cortical regions [4, 5, 7, 36, 37, 46, 47]. In line with this approach, our results may also be applicable to other recurrent cortical circuits that exhibit rich temporal dynamics (e.g., decision-making dynamics in prefrontal cortex [47], temporally structured memories, etc.).

In summary, our results support the view that knowing only the structure of neuronal networks is not sufficient to explain their dynamics [48, 49]. We extend current understanding of the effects of neuromodulation [13, 17, 20, 48] and show that it is possible to control a recurrent neuronal network’s computations without changing its architecture. We found that modulating only neuronal responsiveness enables flexible control of cortical activity. We were also able to combine previously learned modulation states to generate new desired activity patterns, and we demonstrated that employing gain modulation allows one to smoothly and accurately control the duration of network outputs. Our results thus suggest the possibility that gain modulation is a central part of neuronal motor control.

## ACKNOWLEDGEMENTS

We thank the members of the Vogels lab (particularly Everton J. Agnes, Rui P. Costa, William F. Podlaski, and Friedemann Zenke) for their insightful comments and Yayoi Teramoto Kimura for creating the monkey illustration. We also thank Omri Barak, Tim E. Behrens, Rafal Bogacz, Mehrdad Jazayeri, and Lee Susman for their helpful comments. Our work was supported by grants from the Wellcome Trust (TPV and JPS through WT100000, and GH through 202111/Z/16/Z) and the Engineering and Physical Sciences Research Council through the Life Sciences Interface Doctoral Training Centre (University of Oxford) (EP/F500394/1) (JPS).

## AUTHOR CONTRIBUTIONS

JPS, GH, and TPV conceived the study and developed the model. JPS performed simulations for Figs. 1–5 and Supplementary Figs. 1–4, JPS and GH performed simulations for Figs. 6–8 and Supplementary Figs. 5–8. JPS analyzed the results, produced the figures, and wrote the first draft of the manuscript. All authors discussed and iterated on the analysis and its results, and all authors revised the final manuscript.

## DATA AND MATERIALS AVAILABILITY

Sample MATLAB code and other materials are available at https://senselab.med.yale.edu/modeldb. Questions or additional requests should be addressed to JPS.

## 1 METHODS

Our model is specified by a differential equation governing the neuronal firing rates (Eqn. (1)), the gain function Eqn. (2), a set of readout weights, and each neuron’s gain. In the following, we describe our model precisely.

### 1.1 Neuronal dynamics

We model neuronal activity according to Eqn. (1), which we integrate using the ODE45 function (using default parameters) in MATLAB. We do not explicitly model dynamics prior to movement execution; all of our simulations begin at the time of movement onset [4, 11] (except when we use a ramping input in Fig. 2). We choose the initial condition *x*_0_ among the ‘most observable’ modes of the system (i.e., those that elicit the strongest transient dynamics [11]). Specifically, we first linearize the dynamics around its unique equilibrium point *x* = 0 using unit gains (i.e., *g_i_* = 1 for all *i*), and we compute the observability Gramian (a symmetric positive-definite matrix *Q*) of the linearized system [11]. The most observable modes are the top eigenvectors of *Q*. Unless we state otherwise, we choose the eigenvector associated with the largest eigenvalue of *Q* (note that all of its eigenvalues are real and positive) as the initial condition *x*_0_ for the neuronal activity. Following [11], we also scale *x*_0_ so that 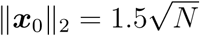.

### 1.2 Biophysical interpretation of Eqn. (1)

Equation (1), together with Eqn. (2), describes how we model neuronal firing rates relative to a baseline rate *r*_0_. In this section, we clarify that one can obtain identical neuronal activity by using a strictly positive gain function *f* and including a constant input *h* in Eqn. (1). Specifically, given a desired baseline firing rate *r*_0_, one can model the neuronal activity as

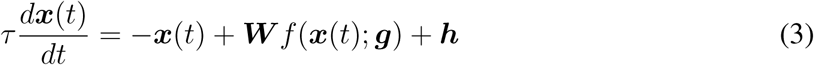

for the same initial condition *x*_0_ that we described above, where *h_i_* = −*r*_0_ ∑_*j*_ *W_ij_* and

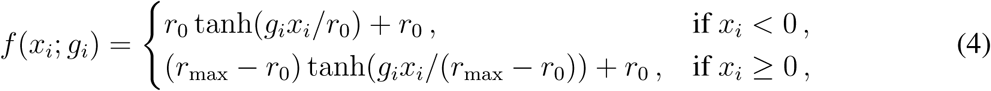

where *r*_max_ is the maximum firing rate. Note that the constant term *h* in Eqn. (4) is necessary to balance the additional *r*_0_ term in Eqn. (4).

### 1.3 Construction of the network architecture

Prior to stability optimization, we generate synaptic weight matrices *W* as detailed in Ref. [11]. In keeping with Dale’s law, these matrices consist of *M* positive (excitatory) columns and *M* negative (inhibitory) columns. We begin with a set of sparse (such that the connection probability between any two neurons is small) and strong weights with nonzero elements set to 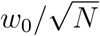 (excitatory) and 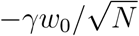 (inhibitory), where 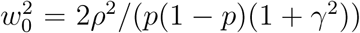 and the connection probability between each two neurons is homogeneous and is given by *p* = 0.1. This construction results in ***W*** having an approximately circular spectrum (i.e., set of eigenvalues) of radius *ρ* (which we set to *ρ* = 10), leading to linear instability before stability optimization (see below). As in Ref. [11], we set the inhibition/excitation ratio *γ* to be *γ* = 3.

After constructing the initial ***W***, we never change any of the excitatory connections. Following [11], we refine the inhibitory connections to minimize an upper bound of ***W***’s ‘spectral abscissa’ (SA) (i.e., the largest real part among the eigenvalues of ***W***) [11]. Briefly, we iteratively update inhibitory weights to follow the negative gradient of this upper bound to the SA. First, the inhibitory weights remain inhibitory (i.e., negative). Second, we maintain a constant ratio (of *γ* = 3) of mean inhibitory weights to mean excitatory weights. Third, we restrict the density of inhibitory connections to be less than or equal to 0.4 to maintain sufficiently sparse connectivity. We observed that this constrained gradient descent usually converges within a few hundred iterations. As was noted in Ref. [11], the SA typically decreases during optimization from 10 to about 0.15. For additional details, see the supplemental information of Ref. [11].

As a proof of principle, we also construct a ‘chaotic’ variant of our neuronal network model (see Fig. 2). These networks are chaotic in the sense that the neuronal dynamics in Eqn. (1) have a positive maximum Lyapunov exponent [50]. We use a synaptic weight matrix ***W*** (as described above) prior to optimization, but now use parameter values of *γ* = 1 and *ρ* = 1.5. We also set *τ* = 20 ms, and we choose the initial condition for each neuron’s activity from a uniform distribution on the interval [−10, 10]. We use only the first 0.5 s of neuronal activity for our simulations of the chaotic network model.

### 1.4 Creating target muscle activity

We generate target muscle activities of duration *t*_tot_ = 500 ms (see Figs. 1–5) and *t*_tot_ = 2, 500 ms (see Figs. 6–8). In each case, we draw muscle activity from a Gaussian process with a covariance function *K* ∈ [0, *t*_tot_] × [0, *t*_tot_] → ℝ_≥0_ that consists of a product of a squared-exponential kernel (to enforce temporal smoothness) and a non-stationary kernel that produces a temporal envelope similar to that of real electromyogram (EMG) data during reaching [4]. Specifically,

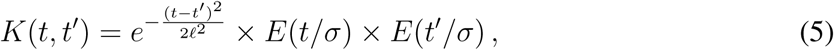

where *E*(*t*) = *te*^(*−t*^2^/4)^. We set *σ* = 110 ms and *ℓ* = 50 ms for movements that last 500 ms and *σ* = 550 ms and *ℓ* = 250 ms for movements that last 2, 500 ms. We also multiply the resulting muscle activity by a scalar to ensure that it has the same order of magnitude as the neuronal activity. We use a sampling rate of 400 Hz for fast movements and 200 Hz for slow movements.

### 1.5 Network output

We compute the network output *z*(*t*) as a weighted linear combination of excitatory neuronal firing rates:

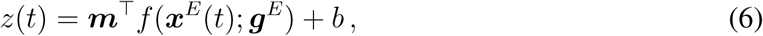

where ***m, x***^*E*^(*t*), ***g***^*E*^ ∈ ℝ^*M*^, the quantity ***x***^*E*^(*t*) is the excitatory neuronal activity, and *M* is the number of excitatory neurons. To ensure that the network output corresponds to realistic muscle activity (see Methods Section 1.4) prior to any training of the neuronal gains, we fit the readout weights ***m*** and the offset *b* to an initial output activity (see Methods Section 1.4) using least-squares regression. To ameliorate any issues of overfitting, we use 100 noisy trials, in which we add white Gaussian noise to the initial condition ***x***_0_ for each trial with a signal-to-noise ratio of 30 dB [11]. Subsequently, the readout weights remain fixed throughout training of the neuronal gains. See our simulation details for each figure for additional details.

### 1.6 Measuring error in network output

We compute the error *∊* between the network output *z* ∈ ℝ^*t*_tot_^ and a target *y* ∈ ℝ^*t*_tot_^ by discretizing time and calculating

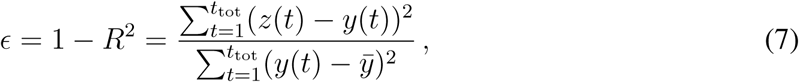

where 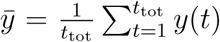 and *R*^2^ is the coefficient of determination (which is often called simply ‘R-squared’). Therefore, an error of *∊* = 1 implies that the performance is as bad as if the output *z* were equal to the mean of the target *y* and thus does not capture any variations in output. When we use multiple readout units, we take the mean error *∊* across all outputs. We use this definition of error throughout the entire paper.

### 1.7 A learning rule for neuronal input–output gains

We devise a reward-based node-perturbation learning rule that is biologically plausible in the sense that it includes only local information and a single scalar reward signal that reflects a system’s recent performance [29, 30]. Our learning rule progressively reduces the error (on average) between the network output and a target output over training iterations. We update the gain *g_i_* for neuron *i* after each training iteration *t_n_* (with *n* = 1, 2, 3, …) according to the following learning rule:

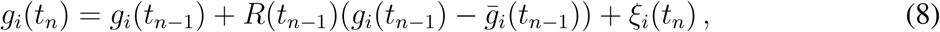

where

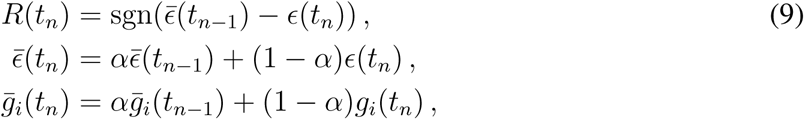

where *∊*(*t_n_*) represents the output error at iteration *t_n_* (see Methods Section 1.6), sgn is the sign function, 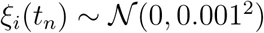 is a Gaussian random variable with mean 0 and standard deviation 0.001, and *α* = 0.3. The initial modulatory signal is *R*(*t*_0_) = 0, and the other initial conditions are 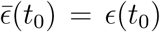 (where *∊*(*t*_0_) is the initial error before training) and 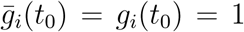. One can interpret the terms 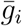 and 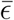 as low-pass-filtered gains and errors, respectively, over recent iterations, with a history controlled by the decay rate *α* [32]. We use these parameter values in all of our simulations in this paper. We find that varying the standard deviation of the noise term *ξ* or the factor *α* has little effect on the learning dynamics (not shown), in line with Ref. [31].

Although our learning rule in Eqn. (8) is similar to reward-modulated ‘exploratory Hebbian’ (EH) synaptic plasticity rules [30–32], we investigate changes in neuronal gains (i.e., the responsiveness of neurons) inside a recurrent neuronal network, rather than synaptic weight changes. The above notwithstanding, we expect our learning rule to perform well for a variety of learning problems. For example, it can solve credit-assignment problems, because one can formulate any such a node-perturbation learning rule as reinforcement learning with a scalar reward [51].

The modulatory signal *R* does not provide information about the sign and magnitude of the error, and it also does not indicate the amount that each readout (if using multiple readouts) contributes to a recent change in performance. The modulatory signal *R* indicates only whether performance is better or worse, on average, compared with previous trials. One can view the modulatory signal as an abstract model for phasic output of dopaminergic systems in the brain [19, 23, 24, 52].

We use the following procedure for updating neuronal gains. We update the gains for iteration *t*_1_ according to Eqn. (8), and we obtain the network output from the gain pattern *g*(*t*_1_). We then calculate the error *∊*(*t*_1_) from the output, and we subsequently calculate the modulatory signal *R*(*t*_1_) and the quantities 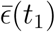 and 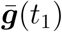 using Eqn. (9). We then repeat this process for all subsequent iterations. If any gain values become negative, we set these to 0. However, this happened very rarely in our computations, and we observed it only when we used 60,000 training iterations (i.e., in Figs. 3e and 6b).

### 1.8 Alternative learning rule

One can also adapt our learning rule so that learning ceases when the reward signal *R*(*t_n_*) saturates at a sufficiently small value. A way to achieve this is by instead placing the noise term *ξ_i_* inside the brackets in Eqn. (8), so that the modulatory signal *R* multiplies *ξ_i_*, together with changing the sgn function in Eqn. (9) to the tanh function. This yields the following learning rule:

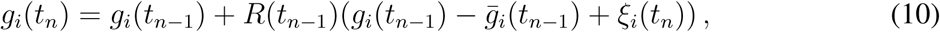

where

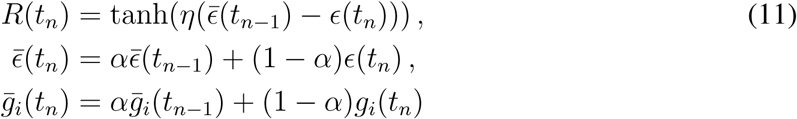

and *η* = 50,000 controls the slope of the tanh function at 0 (i.e., when the low-pass-filtered error 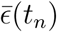 matches the current error *∊*(*t_n_*)). Learning now stops when 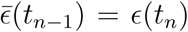; see the purple curve in Fig. 2a. We achieve a qualitatively similar learning performance by using Eqns. (10) and (11) instead of Eqns. (8) and (9), respectively. (Compare the purple and red curves in Fig. 2a.)

### 1.9 Generating groups for group-based gain modulation

For coarse-grained (i.e., grouped) gain modulation, we generate *n* (modulatory) groups, and we independently modulate each group using one external ‘modulatory unit’. Our generation mechanism for random groups is as follows. For each of the *n* groups, we choose *N/n* neurons (where *N* is the total number of neurons in the network) uniformly at random without replacement. If *n* does not divide *N*, we assign the remaining neurons to groups uniformly at random.

When using specialized groupings (see Figs. 3b,c and Supplementary Figs. 2a–d) for a particular target movement, we obtain groups by applying *k*-means clustering (where *k* is the desired number of groups) to 10 gain patterns that we obtain from 10 prior independent training sessions (using neuron-specific control) on the same target and which correspond to the minimum error for each training session. We thus apply *k*-means clustering to a matrix of size *N* × 10, where row *i* has the gain values for neuron *i* from the 10 independent training sessions to the same target. Applying *k*-means clustering then generates groupings in which neurons in the same group tend to have similar gain values following training using neuron-specific modulation.

### 1.10 Simulation details

We now give details about our simulations for Figures 1–8. For further information, see our full simulation details in the Supplementary Information section of this file and see our Supplementary Maths Note for our mathematical derivations.

**Figure 1.** We simulate two different electromyograms (EMGs) (see Methods Section 1.4) of muscle activities (initial reach and target reach) that each last 0.5 s (see Figs. 1a,f). We use a network of *N* = 200 neurons and sample transient neuronal firing rates that last 0.5 s following the initial condition *x*_0_ of the neuronal activity (see Methods Section 1.1). We fit the readout weights over 100 trials, in which we add white Gaussian noise to the initial condition *x*_0_ (with a signal-to-noise ratio of 30 dB) using least-squares regression so that the network output, with all gains set to 1, generates the initial reach (see Methods Section 1.5). We use the same readout weights throughout all training, and we use only one readout unit for each simulation.

For each training iteration of the neuronal gains (to generate a target movement), we use the initial condition *x*_0_ at time *t* = 0 (see Methods Section 1.1). We calculate the subsequent network output as described in Methods Section 1.5, and we update the neuronal gains according to Eqn. (8). We repeat this process for 18,000 training iterations (which corresponds to 2.5 hours of training time), which is enough training time for the error to saturate (see Fig. 1d). We run 10 independent training sessions on the same target, and we plot these results in Figs. 1d,e.

**Figure 2.** We train neuronal gains on the same task as the one that we showed in Fig. 1d using 3 alternative models. For one model, we use a ramping input to the neuronal activity in Eqn. (1) as a model of preparatory activity prior to movement onset [4, 11]. We use the same ramping input function as the one that was used in Ref. [11]. It is exp(*t/τ*_on_) for *t* < 0 s and exp(*−t/τ*_off_) after movement onset (*t* ≥ 0), with an onset time of *τ*_on_ = 400 ms and an offset time of *τ*_off_ = 2 ms. Gain changes that result from learning now also affect the neuronal activity at *t* = 0 (i.e., at movement onset).

We also train a ‘chaotic’ [34] variant of our model (see Methods Section 1.3, where we describe how we construct such a model), and we use the first 0.5 s of neuronal activity.

Finally, we use an alternative learning rule (see Eqns. (10) and (11)) in which learning stops automatically when the difference between network outputs in successive training iterations becomes sufficiently small (see Methods Section 1.7).

**Figure 3.** For Figs. 3b,c, we generate 5 different target outputs and run 10 independent training sessions for each target. For the random groupings (see Methods Section 1.9), we use different independently generated random groups for each simulation. For the specialized groups (see Methods Section 1.9), for a given number of groups, we use the same grouping in all simulations.

We now explain how we determine specialized groups that are shared by multiple movements (i.e., we use the same grouping for learning multiple movements); see the plots in Fig. 3c and Supplementary Figs. 2b–d. We apply *k*-means clustering (where *k* is the desired number of groups) across all of the gain patterns that we obtain using neuron-specific modulation for each of the movements.

For the task that we just described above, we consider various different numbers of groups (using random groupings) for networks with *N* = 100, *N* = 200, and *N* = 400 neurons. We again perform 10 independent training sessions for each network, target, and number of groups. We fit the readout weights so that each scenario generates the same network output when all gains are set to 1. The readout weights remain fixed throughout training. We plot these results in Fig. 3d and Supplementary Figs. 2e–h.

When we use multiple readout units, we generate 10 different initial and target outputs for each readout unit. We run independent training sessions for these 10 sets of target outputs and calculate mean errors across the 10 training sessions. For a given number of readout units, we use the same sets of initial and target outputs for all 3 network sizes and each number of random modulatory groups. We thus fit readout weights so that each scenario generates the same output with all gains set to 1. The readout weights remain fixed throughout training. We use 60,000 (instead of 18,000) training iterations to ensure error saturation.

**Figure 4.** To create libraries of learned movements, we train a network of 400 neurons and 40 random groups (see Methods Section 1.9) on each of 100 different target movements independently. (In other words, this generates 100 different gain patterns, with one for each movement.) For library sizes of *l* ∈ {1, 2, …,50}, we choose 100 samples of *l* movements (from the learned gain patterns and their outputs) uniformly at random without replacement for each *l*. We then fit the set of movements in each of the 100 sample libraries using least-squares regression for each of 100 hitherto-untrained novel target movements. We constrain the fitting coefficients *c_j_* from the least-squares regression by requiring that *c_j_* ≥ 0 for all *j* and 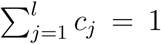. We calculate the fit error (i.e., the error between the fit and the target), the output error (i.e., the error between the output and the target), and the error between the fit and the output for each of the 100 novel target movements, each of the 100 library samples, and each *l*. In Fig. 4, we show results for up to *l* = 20 library elements, whereas in Supplementary Fig. 3, we show results for up to *l* = 50 library elements.

**Figure 5.** We train the same 200-neuron weight matrix that we used in Fig. 1 on the same task as the one that we showed in Figs. 1d–f, except with a baseline rate of *r*_0_ = 5 Hz in Eqn. (2). We also repeat the simulations that we performed in Fig. 4 for the baseline rate *r*_0_ = 5 Hz.

**Figure 6.** In each of these simulations, we use a network of 400 neurons and 40 random modulatory groups (see Methods Section 1.9). We construct ‘slow’ (2.5 s) target movements with *σ* = 550 ms and *ℓ* = 250 ms in Eqn. (5). We then construct a ‘fast’ (0.5 s) variant of each movement. Each movement variant has 500 evenly-spaced points, and we augment 400 instances of 0 values to the final 2 s of the movement to ensure that both movement variants have the same length.

Note that we are modelling network output as a proxy for muscle-force activity. When we study whether we can generate the same movement that lasts 5 times longer (see Figs. 6–8), we scale the duration of the muscle activity without changing its amplitude. To actually generate the same movement so that it lasts 5 times longer, we also need to scale the amplitude of the muscle activity by the factor 1/5^2^ = 1/25. To demonstrate the effectiveness of learning through gain modulation, we omit this scaling, so the tasks on which we train are more difficult ones, as the target activity without the scaling has a substantially larger amplitude throughout the movement. However, we find that learning through gain modulation can also account for this scaling of muscle activity when performing movements at different speeds (see Supplementary Fig. 8). Alternatively, it may be possible for gain modulation of downstream motoneurons in the spinal cord to account for scaling of the amplitude of muscle activity when performing movements at different speeds (for example, see Ref. [21]).

For Fig. 6b, we fit readout weights using least-squares regression, such that with all gains set to 1, the network output approximates the fast variant. We then train gain patterns using our learning rule in Eqns. (8) and (9) so that the network output generates the slow-movement variant. (The initial condition *x*_0_ and readout weights remain fixed.) We use 60, 000 training iterations, and we run 10 independent training sessions for each of 10 different target movements.

For Fig. 6c, we perform the task that we described in the paragraph above using a gradient-descent training procedure with gradients that we obtain from back-propagation [53]. Together with learning the gain pattern for the slow variant, we jointly optimize a single set of readout weights (shared by both the fast-movement and slow-movement variants) as part of the same training procedure. The gains are still fixed at 1 for the fast variant. The cost function for the training procedure is equal to the squared Euclidean 2-norm between actual network outputs (fast and slow) and target outputs (fast and slow) plus the Euclidean 2-norm of the readout weights, where the latter acts as a regularizer. We run gradient descent for 500 iterations, which is well after the cost has stopped decreasing.

For each of the 10 trained movements that we described earlier in this section, we extract the mean minimum error across all simulations for both the outputs obtained via our learning rule (see Supplementary Fig. 5a) and the outputs obtained via back-propagation (see Supplementary Fig. 5b). We then linearly interpolate between the learned gain patterns for the fast and slow outputs, and we calculate the error between the output and the target movement at the interpolated speed. (See the top panel of Fig. 6d.)

For Figs. 6d–f, we train networks to generate a pair of target movements in response to a corresponding pair of orthogonal initial conditions at fast and slow speeds and also at each of 5 intermediate, evenly-spaced speeds in between these extremes. To do this, we parametrize the gain pattern of speed index *s* (with *s* ∈ {1, …,7}) as a convex combination of a gain pattern ***g***_*s*=1_ for fast movements and a gain pattern ***g***_*s*=7_ for slow movements, with interpolation coefficients of *λ_s_* (with ***g***_*s*_ = *λ_s_****g***_*s*=1_ + (1 − *λ_s_*)***g***_*s*=7_, *λ*_1_ = 1, and *λ*_7_ = 0). We optimize (using back-propagation, as discussed above) over ***g***_*s*=1_, ***g***_*s*=7_, the 5 interpolation coefficients *λ_s_* (with *s* ∈ {2, …,6}), and a single set of readout weights. For a given speed *s*, we use the gain pattern ***g***_*s*_ for both movements. We call the collection of gain patterns ***g***_*s*_ for *s* ∈ {1, …,7} the gain manifold for speed control (or the ‘speed manifold’, as a shorthand).

**Figure 7.** We train (using back-propagation, as we discussed previously) a 400-neuron network with 40 random modulatory groups (see Methods Section 1.9) to generate each of 10 different movement shapes at 7 different, evenly-spaced speeds (ranging from the fast variant to the slow variant) using a fixed initial condition *x*_0_. To jointly learn gain patterns that control movement shape and speed, we parametrize each gain pattern as the element-wise product of a gain pattern that encodes shape (which we use at each speed for a given shape) and a gain pattern that encodes speed (which we use at each shape for a given speed). We again parametrize (see our simulation details for Fig. 6) the gain pattern that encodes speed index *s* (with *s* ∈ {1, …,7}) as a convex combination of two common endpoints, ***g***_*s*=1_ (which we use for the fast-movement variants) and ***g***_*s*=7_ (which we use for the slow-movement variants). (We call this the ‘speed manifold’; see our simulation details for Fig. 6.) We thus optimize over 10 gain patterns for movement shape, 2 gain patterns each for fast and slow movement speeds, 5 speed-interpolation coefficients, and a single set of readout weights.

In Fig. 7c, we calculate the mean error between the network output and the target over the 10 target movements when generating gain patterns for movement speed by linearly interpolating between the trained fast (***g***_*s*=1_) and slow (***g***_*s*=7_) gain patterns.

**Figure 8.** We use the 10 trained gain patterns for movement shapes, as well as the speed manifold from Fig. 7 (see our simulation details for Fig. 7). Using our learning rule from Eqns. (8) and (9), we train the 10 coefficients *c*_1_, …, *c*_10_ (see Fig. 8a) to construct a new gain pattern from the 10 existing shape-specific gain patterns that, together with the speed manifold, generates a new target movement at the fast and slow speeds. Specifically, we replace the gains *g_i_* (for *i* ∈ {1, …, *N*}) with the coefficients *c_i_* (for *i* ∈ {1, …,10}) in Eqns. (8) and (9). We use the mean of the errors at the fast and slow speeds in our learning rule. To generate the network output at the fast and slow speeds, respectively, we calculate the element-wise product between the newly-constructed gain pattern and the fast and slow gain pattern, respectively, on the speed manifold. We independently train, using 10,000 training iterations, the coefficients *c*_1_, …, *c*_10_ on each of the 100 target movements that we used for Fig. 4. As a control, we calculate the mean error between the network output and the target over the 100 target movements when choosing one of the 100 newly-learned gain patterns uniformly at random without replacement. (See the grey curve in Fig. 8c.)

Additionally, instead of learning to combine gain patterns using the method that we described in the previous paragraph, we determine coefficients *c*_1_, …, *c*_10_ using a least-squares regression by fitting the 10 learned movements to each of the 100 target movements at the fast and slow speeds simultaneously and requiring that *c_j_* ≥ 0 for all *j* and 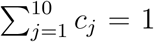. (See the black dashed curve in Fig. 8c.)

Finally, we plot the Pearson correlation coefficient between pairs of target movements versus the Pearson correlation coefficient between corresponding pairs of learned coefficients *c*_1_, …, *c*_10_. Note that we are unlikely to observe correlation values close to −1 between pairs of combination coefficients because the coefficients *c*_1_, …, *c*_10_ are likely to form a convex combination, as in Fig. 4 (in fact we calculate the mean sum of the coefficients to be approximately 0.91). In our visualization, we plot only 1,000 of the 4,950 data points. (We choose these points uniformly at random.)

## SUPPLEMENTARY INFORMATION

### Supplementary maths note

#### Analysis of the effects of identically changing the gain of all neurons

To examine the effects of gain modulation on neuronal dynamics when identically changing all neuronal gains (i.e., *g_i_* = *g* for all *i*), we construct a Taylor expansion of *f*(*x_i_; g_i_*) from Eqn. (2) around *x* = 0. By keeping only leading-order terms, we obtain *f*(*x_i_*;*g*) ≈ *gx_i_*, and substituting this expression into Eqn. (1) yields 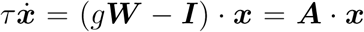, where ***I*** is the identity matrix and ***A*** = *g****W*** − ***I***. Empirically, we find this linear approximation to be valid in a large basin of attraction around the equilibrium point.

Changing the gain from *g* to *g*′ multiplies the imaginary part of the spectrum of ***A*** by the factor *g′/g*. (Subtracting the identity matrix does not affect the imaginary part of the spectrum of ***A***.) This, in turn, multiplies the frequency of the associated solution of the linearized dynamics of *x*(*t*) by the factor of *g′/g*.

A change in gain also causes changes in the real parts of the eigenvalues of ***A***. Specifically, increasing the gain causes the real parts of all but one of the eigenvalues of *g**W*** to increase (i.e., the eigenvalues of ***A*** get closer to the imaginary axis), generally causing a slower decay of activity towards the equilibrium [54]. The real part of the remaining eigenvalue, which is associated with the eigenvector 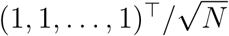 (see Ref. [11]), becomes more negative with increasing gain, resulting in faster decay of the neuronal dynamics. However, this effect is small in comparison with the slowing of the decay due to the changes of the real parts of all of the other eigenvalues.

#### Analysis of linear combinations of gain patterns and their associated neuronal dynamics

In Fig. 4 and Supplementary Figs. 3 and 4, we illustrated that there is a consistent mapping between learned gain patterns and their outputs. Specifically, we illustrated that for a library of *l* gain patterns (***g***_1_, …, ***g***_*l*_), a convex combination *c*_1_*F*(***g***_1_) + … + *c_l_F*(***g***_*l*_) (so *c_j_* ≥ 0 for all *j* and 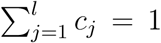) of their corresponding outputs (which we denote by *F*) approximates the output *F*(*c*_1_***g***_1_ + … + *c_l_****g***_*l*_) that we obtain by combining the gain patterns with the same coefficients (see Fig. 4). Note that the subscript index *j* denotes the library element *j* and is not a neuron index. We now provide some mathematical understanding of this approximation by studying linearized solutions of the neuronal dynamics. Because the network output is a linear combination of the neuronal firing rates, it is sufficient to study convex combinations of internal neuronal activity *x* directly, rather than convex combinations of network outputs.

For a convex combination (i.e., a weighted mean) of *l* vectors or matrices *ϕ* with weights *c_j_*, it is convenient to use the following notation:

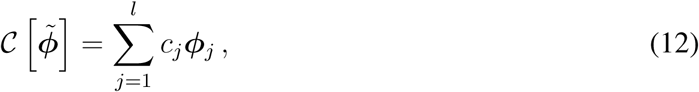

where the tilde in the square brackets is a reminder that we are summing over the index of the associated library terms. When we employ matrix multiplication in the following analysis, it is convenient to represent a gain pattern ***g***_*j*_ ∈ ℝ^*N*^ using the matrix notation ***G***_*j*_ = diag(***g***_*j*_) (that is, the neuronal gains are elements along the diagonal of ***G***_*j*_ ∈ ℝ^*N*×*N*^, all other elements are 0, and the index *j* denotes library element *j*). Using this notation, the solution ***x***_*j*_(*t*) ∈ ℝ^*N*^ of the linearized dynamics of Eqn. (1) around ***x*** = 0 is given by

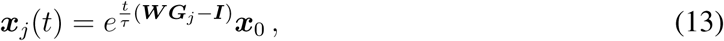

under the assumption that there are *N* distinct eigenvectors for the matrix ***WG***_*j*_ − ***I*** and that we are away from any bifurcations. Let

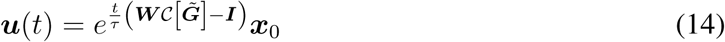

denote the neuronal activity that results from a convex combination 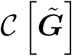 of gain patterns. We need to show that ***u***(*t*) is approximately the same as the convex combination of the individual neuronal dynamics ***x***_*j*_(*t*) with the same coefficients *c_j_*. That is, we need to show that the difference

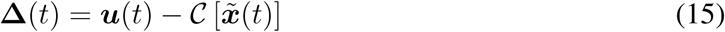

is small with respect to the magnitude of the neuronal activity. We first note that 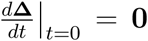, which we prove as follows:

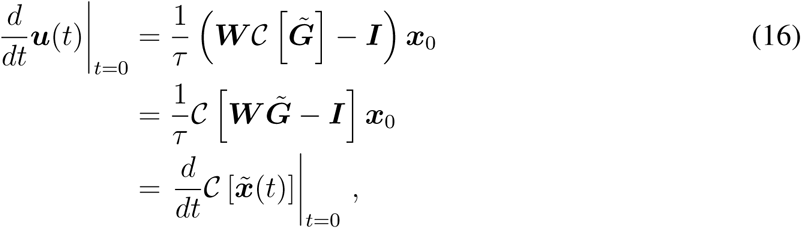

where we used the fact that 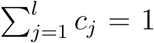 to go from the first to the second line, and we note that the matrices ***W*** and ***I*** do not depend on the gain patterns.

To see whether we can also expect Δ(*t*) to be small for *t* > 0, it is useful to consider the power-series expansion of the matrix exponentials on the right-hand side of Eqn. (15):

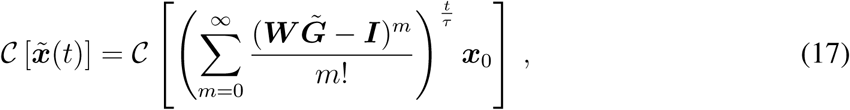

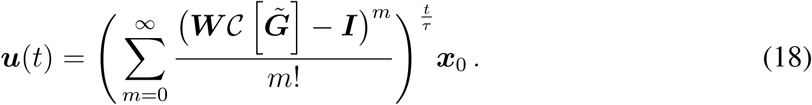

We observe in numerical simulations (not shown) that power-series expansions of this form are accurate descriptions of the associated neuronal dynamics up to second order in *m*. We therefore truncate to *m* = 2, and we evaluate the difference of Eqns. (17) and (18):

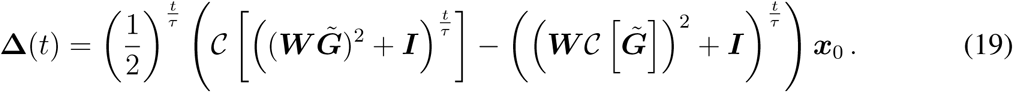

We need to check if the right-hand side of Eqn. (19) is small compared to the neuronal dynamics (i.e., compared to Eqn. (17)). One way to check if this holds at certain times *t* is to substitute values of *t* into Eqns. (19) and (17) and calculate the ratio of the norms of these two expressions. Setting *t* = *τ* — at *t* = *τ* = 200 ms, the neuronal dynamics are close having reached their maximum amplitude (see Supplementary Fig. 2e) — yields

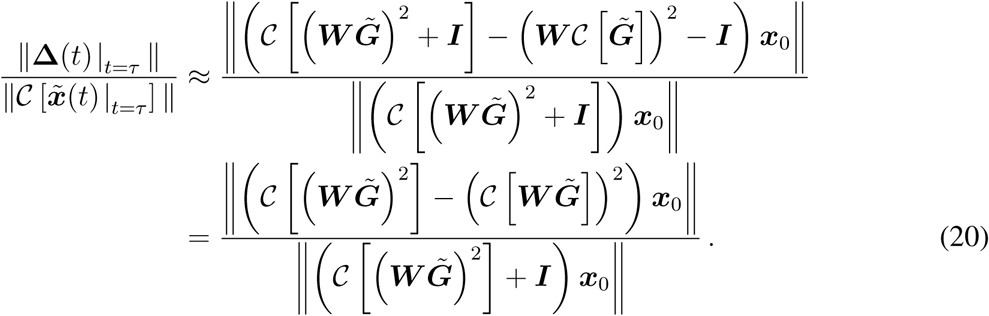

We now study the magnitude of the numerator and the denominator of Eqn. (20) and show that the ratio of the former to the latter is small. Both the numerator and the denominator scale approximately in linear proportion to the norm of the product of ***W***^2^ and ***x***_0_. (The identity matrix in the denominator is small compared to ***W***^2^.) The main difference between the numerator and denominator is their dependencies on the gain patterns ***G***_*j*_. The numerator scales approximately proportionally to a ‘weighted variance’ of the gain patterns, whereas the denominator scales approximately proportionally to a weighted mean of the squared gain patterns. Because our learned gain patterns are typically narrowly distributed, with a mean of 1 and approximate standard deviation of 0.15 (see Supplementary Fig. 3a), this ratio is expected to be small. Numerically, we confirm that the normalized error in Eqn. (20) is indeed small (on the order of 10^−3^) and actually decreases with an increasing library size. This corroborates the results of Fig. 4 of the main text.

Finally, although we restricted our discussion above to a linear gain function, we note that our numerical simulations suggest that Eqn. (15) is also small for the nonlinear gain function of Eqn. (2) (see Fig. 4 and Fig. 5f) that we used throughout the main text.

**Supplementary Figure 1:**
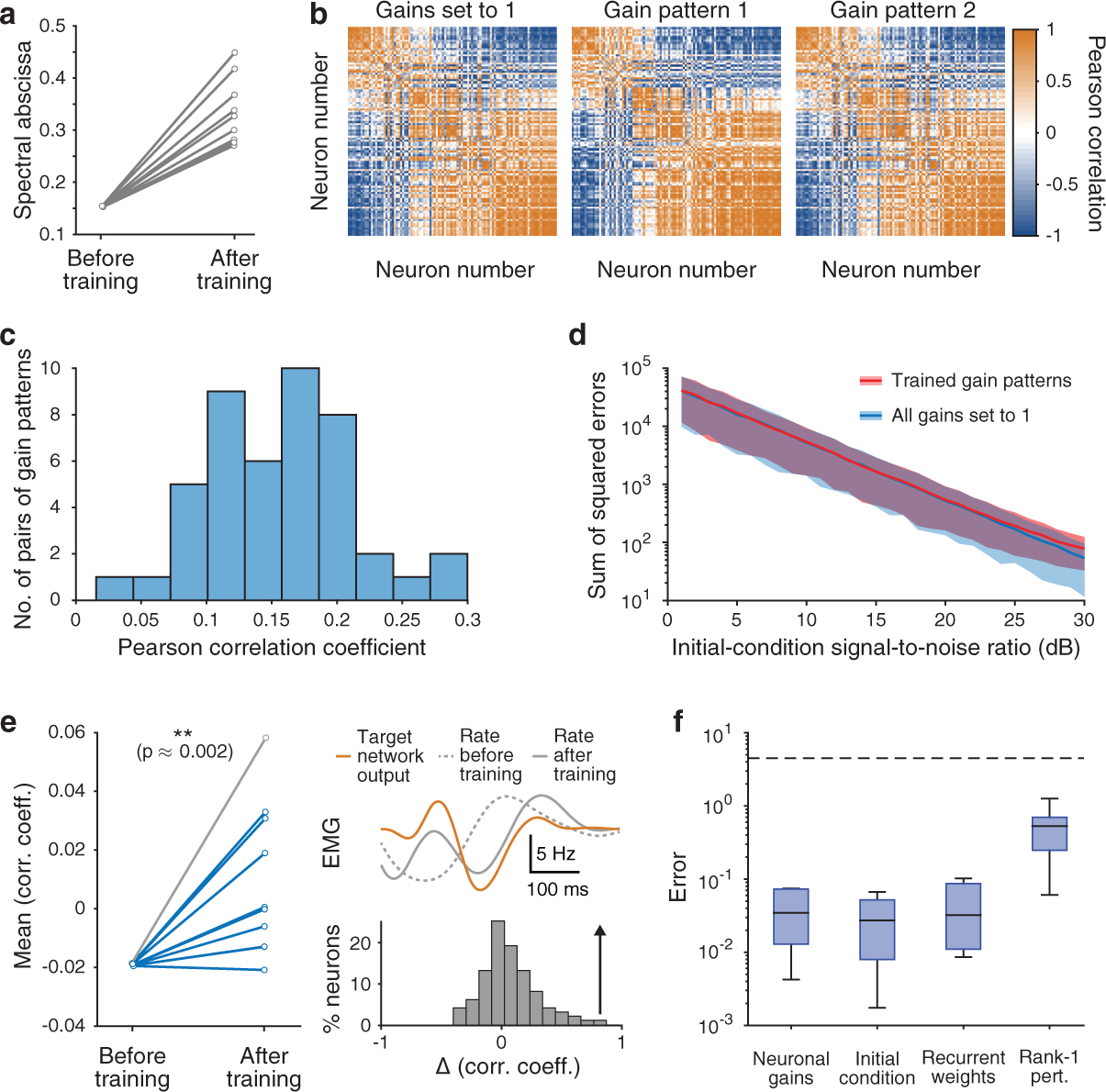
Further effects of neuron-specific gain modulation. (**a**) Changes in the largest real part in the spectrum of ***W*** × diag(***g***) that result from 10 different training sessions (see Methods Section 1.10). Although this change seems substantial, the resulting firing-rate activity does not change dramatically. (For example, see panel (b) and the far left panel of Fig. 2b.) (**b**) Pearson correlation matrices of the firing rates for all pairs of neurons with (left) all gains set to 1 and (centre and right) two (of the ten) trained gain patterns for the task in Fig. 1d. The order of neurons is the same in all three matrices. As a result of training, there is not a substantial reorganisation in Pearson correlations between pairs of neurons. (**c**) Histogram of the Pearson correlation coefficients between the 45 pairs of the 10 trained gain patterns for the task in Fig. 1d. (**d**) Mean error between the network output with white Gaussian noise added to the initial condition ***x***_0_ and the network output without noise added to ***x***_0_ for various signal-to-noise ratios for 1,000 different samples of such noise. We plot results with all gains set to 1 (blue) and the 10 trained gain patterns (red) for the task in Fig. 1d. Shading indicates 1 standard deviation. (See our full simulation details for further information.) (**e**) (Left) The mean Pearson correlation coefficient between the neuronal firing rates and the target increases after training. (We show 10 training sessions and we used a paired Wilcoxon signed rank one-sided test to generate a p-value of *p* ≈ 0.002.) (Bottom right) Example change in Pearson correlation coefficients between the 200 neurons’ firing rates and the target after training for the trial in grey in the left panel. (Top right) Example of a substantial change in the dynamics of one neuron after training. (**f**) Box plots of the errors after training independently on 10 different target movements using back-propagation for four different scenarios. In these examples, we train either the neuronal gains, the initial condition, the recurrent synaptic weight matrix, or a rank-1 perturbation of the synaptic weight matrix (see our full simulation details). The dashed black line is the mean error over the 10 target movements before training. (Centre lines indicate median errors, boxes indicate 25th to 75th percentiles, whiskers indicate ±1.5 × the interquartile range, and dots indicate training sessions whose error values lie outside whiskers.) (We use 200-neuron networks for all simulations in this figure.)

**Supplementary Figure 2:**
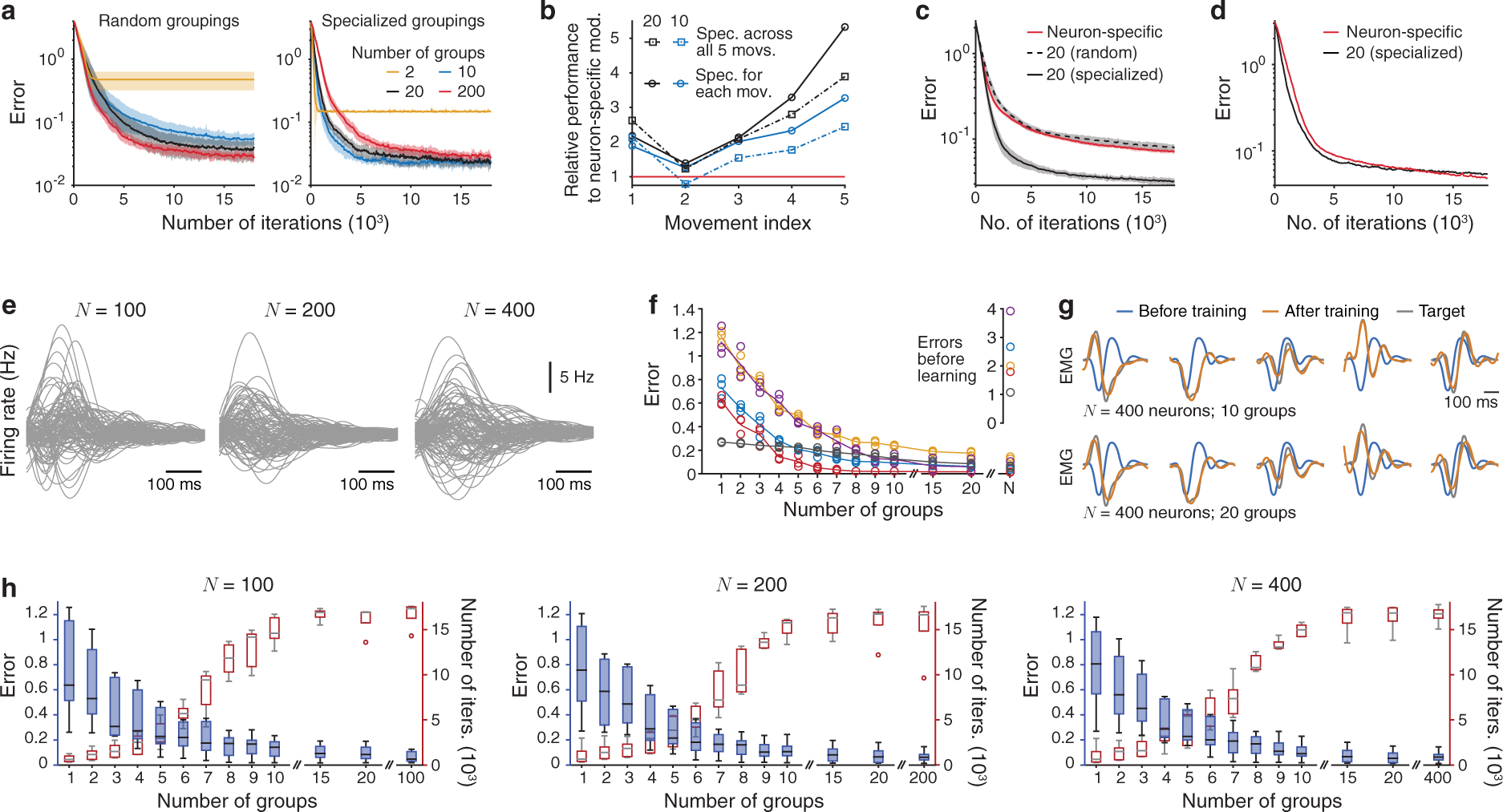
Additional results for grouped gain modulation. (**a**) Mean error over 10 training sessions (where shading indicates one standard deviation) using (left) random and (right) specialized groupings for 2, 10, 20, and 200 (i.e., neuron-specific) groups. (See Methods Sections 1.9 and 1.10.) The target output is the same as in Fig. 1. (**b**) Relative improvement in performance compared with neuron-specific modulation for each of 5 movements when using specialized groups shared across all (squares) or for each (circles) of the 5 movements using either 10 (blue) or 20 (black) groups. A value of 2 implies that the error is 2 times smaller after training compared to neuron-specific modulation. We indicate the performance of neuron-specific modulation using the red line. (See our full simulation details for further information.) (**c**) Mean error over 10 training sessions (where shading indicates one standard deviation) when learning 5 movements using either 20 specialized groups (shared across all 5 movements), 20 random groups, or neuron-specific modulation. (**d**) Mean error over 10 training sessions when learning 10 novel movements using the specialized grouping (with 20 groups) shared across the 5 previously trained movements from panel (c). (**e**) The firing rates of 50 inhibitory and 50 excitatory neurons for each of the three different networks sizes. (**f**) The curves give the mean error over 10 training sessions and across the 3 networks for each of 5 targets. The circles represent the mean error for each network, and the different colours indicate each of 5 different target outputs (see Methods Section 1.10). (**g**) Outputs for all five targets from the trial that produces the median error for the 400-neuron network for the cases of 10 and 20 groups. (**h**) Box plots (in blue) of the minimum error after training for different numbers of groups and the 3 different network sizes. (These are the same data that we plotted in panel (f).) We also include box plots (in red) for the minimum number of iterations required before the error is within 1 % of the minimum error. (Centre lines indicate median errors, boxes indicate 25th to 75th percentiles, whiskers indicate ± 1.5× the interquartile range, and dots indicate training sessions whose error values lie outside whiskers.)

**Supplementary Figure 3:**
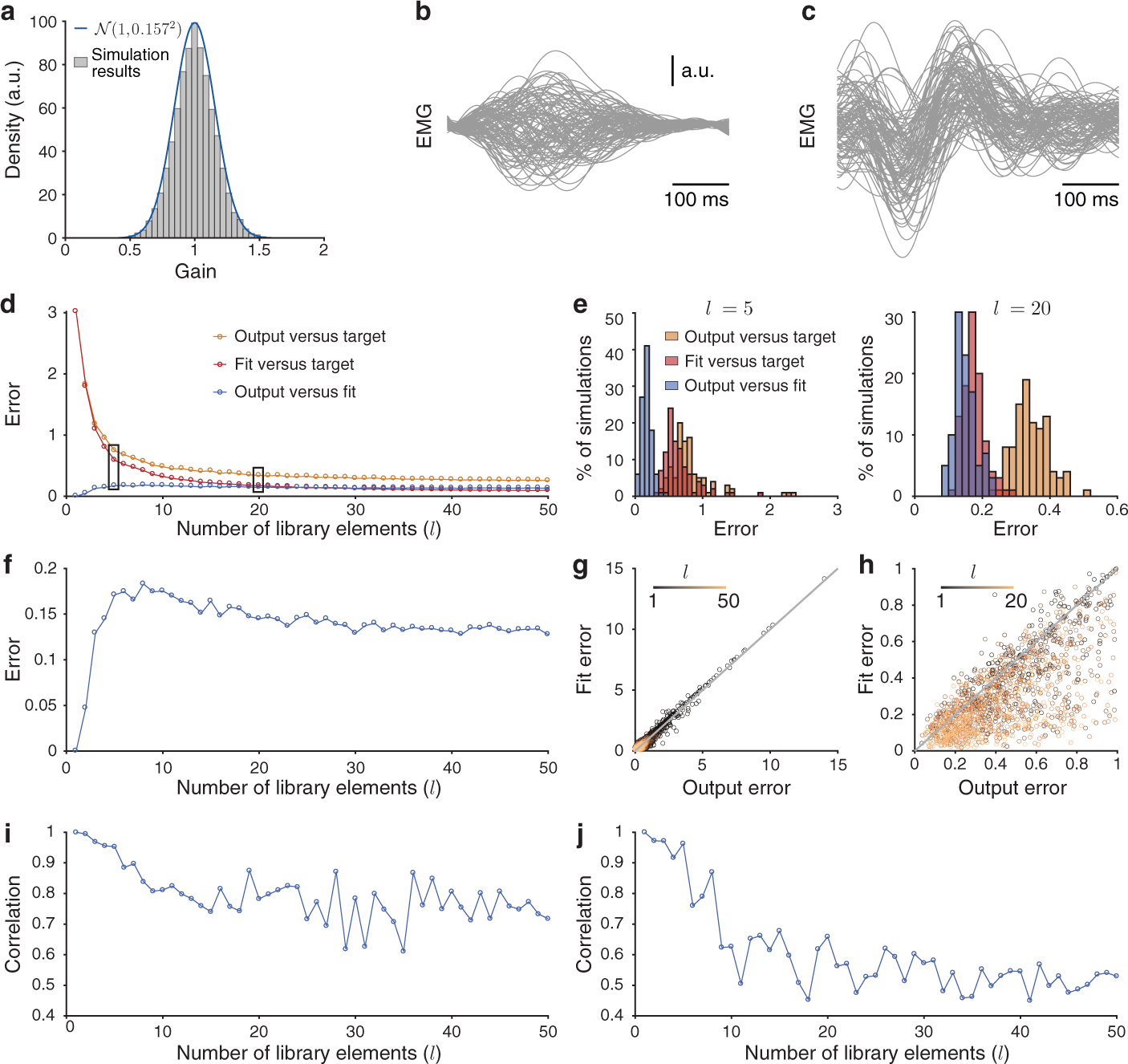
Additional results for gain patterns providing motor primitives. (**a**) The resulting distribution of gains from training independently on each of 100 targets (see Methods Section 1.10). The distribution of the gain patterns resembles a normal distribution (blue curve) with the same mean and variance as those in Fig. 1e. (**b**) Each output from the 100 trained gain patterns. (**c**) Outputs of 100 randomly-generated gain patterns from the distribution in panel (a). (See Methods Section 1.10 and our full simulation details for further details.) The outputs are substantially more homogeneous than those in panel (b) and likely would not constitute a good library for movement generation. (**d**) The same plot as in Fig. 4d, but for up to *l* = 50 library elements. (**e**) The distributions of errors across 100 different libraries for (left)*l* = 5 and (right) *l* = 20. (Note the difference in horizontal-axis scales in the two plots.) (**f**) The error between the output and the fit from panel (d) with a different vertical axis scale. (**g**) The same plot as in Fig. 4c, but for *l* = 1, …,50 and with extended axes. Each point represents the 50th-smallest error between the output and the fit across 100 novel target movements for each of 100 randomly-generated combinations of *l* library elements. We show the identity line in grey. (**h**) The same as in panel (g), but each point represents the 50th-smallest error between the output and the fit across the 100 libraries for each of the 100 novel target movements. We plot these data in the square [0, 1] × [0, 1] and for *l* = 1, …, 20. (**i**) For the data in panel (g), we plot the Pearson correlation coefficient between the output and the fit errors over the 100 randomly-generated libraries for each number of library elements (up to *l* = 50). (**j**) For the data in panel (h), we plot the Pearson correlation coefficient between the output and the fit errors over the 100 novel target movements for each number of library elements (up to *l* = 50).

**Supplementary Figure 4:**
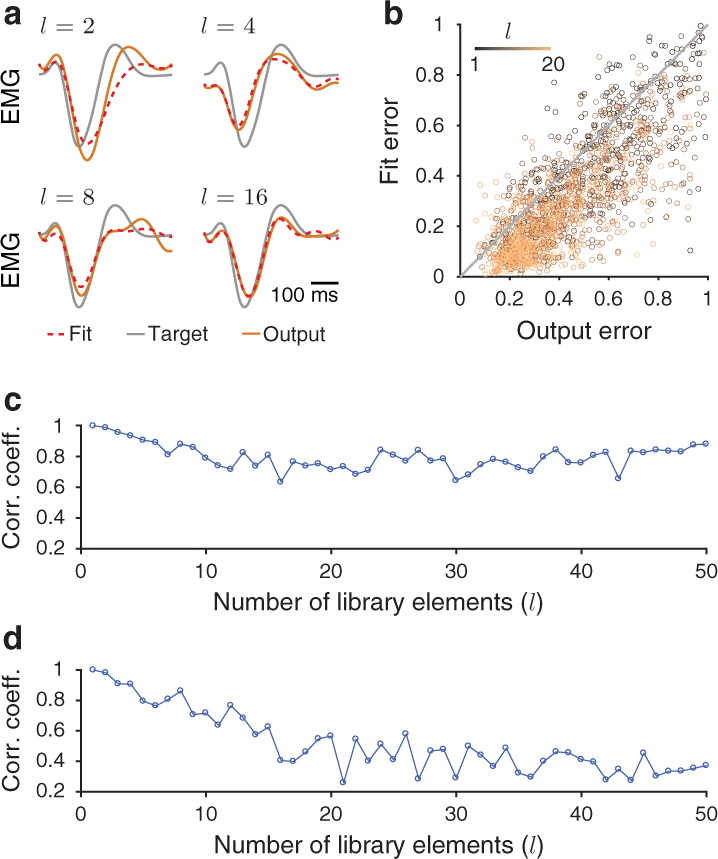
Gain patterns as motor primitives with *r*_0_ = 5 Hz. (**a**) Example target (grey), fit (dashed red), and output (orange) that produces the 50th-smallest output error over 100 randomly-generated combinations (see Methods Section 1.10 for a description of the generation process) of *l* library elements using *l* = 2,*l* = 4,*l* = 8, and *l* = 16. (**b**) Fit error versus the output error for 100 randomly-generated combinations of *l* library elements for *l* = 1, …, 20. Each point represents the 50th-smallest error between the output and the fit across 100 novel target movements. We show the identity line in grey. (**c**) For the data in panel (b), we plot the Pearson correlation coefficient between the output and the fit errors over the 100 randomly-generated combinations of library elements for each number of library elements (up to *l* = 50). (**d**) The same as panel (c), but for data corresponding to the 50th-smallest error for each of the 100 novel target movements, rather than for each randomly-generated combination of library elements (up to *l* = 50) (see our full simulation details for further information). Compare panels (c) and (d) of this figure with panels (i) and (j) in Supplementary Fig. 3.

**Supplementary Figure 5:**
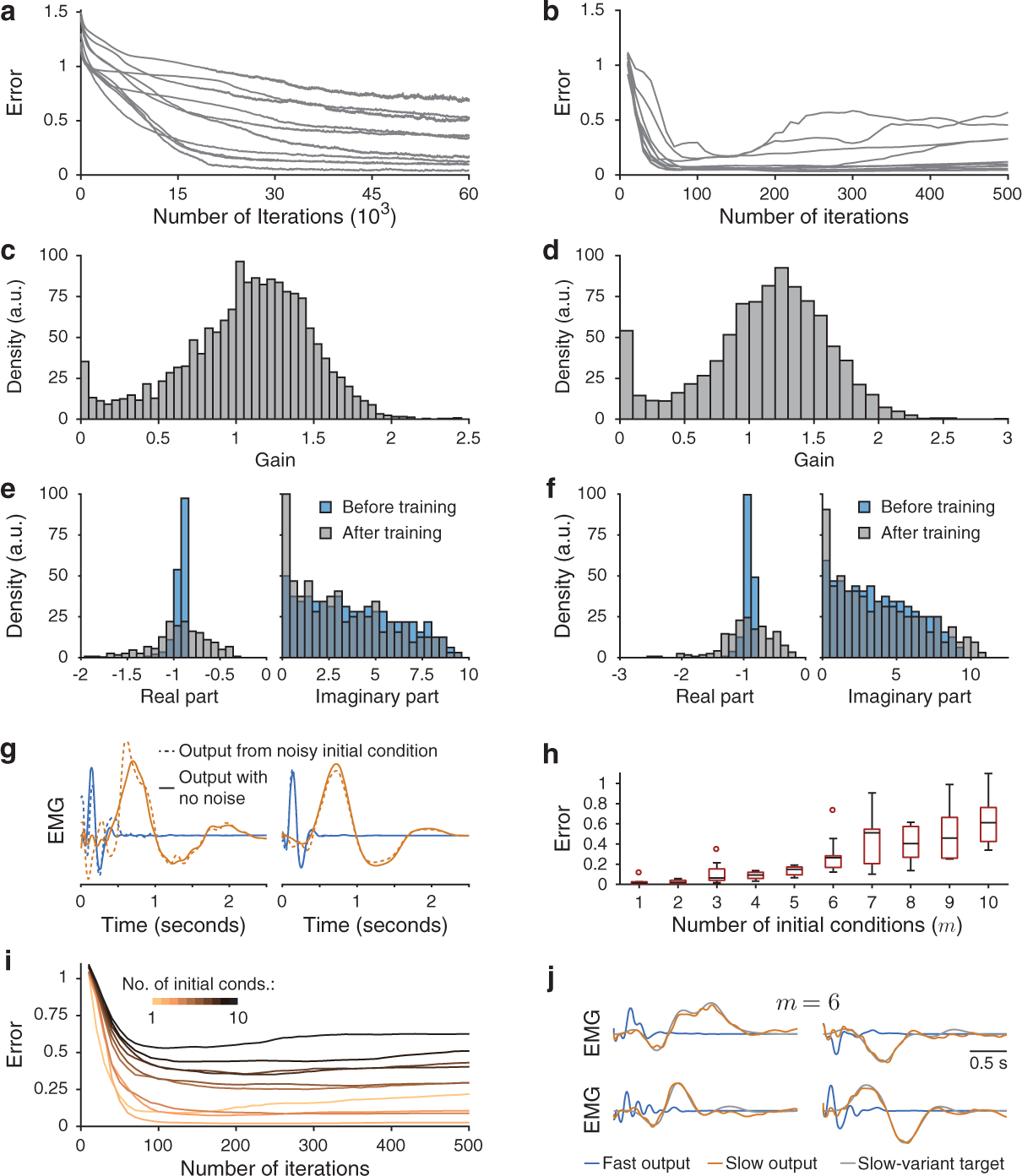
Additional results for controlling movement speeds through gain modulation. (**a**) Mean error over 10 training sessions for each of 10 different movements when learning gain patterns for slow-movement variants using our reward-based learning rule (see Methods Section 1.10). (**b**) Mean error over 10 training sessions for the same 10 movements when instead learning gain patterns for slow-movement variants using a back-propagation algorithm (see Methods Section 1.10). (**c**) Distribution of gains for the slow-movement variants across all training sessions using our reward-based learning rule. (**d**) Distribution of gains for the slow-movement variants across all training sessions when using back-propagation. (**e**) Histograms of the real and imaginary parts of the eigenvalues of the linearization of Eqn. (1) before and after training using our reward-based rule for the example in Fig. 6b. (**f**) Histograms of the real and imaginary parts of the eigenvalues of the linearization of Eqn. (1) before and after training using the back-propagation algorithm for the example in Fig. 6c. (**g**) On the left and right, respectively, we show the same outputs that we plotted in Figs. 6b and 6c, but we now add white Gaussian noise (with a signal-to-noise ratio of 4 dB) to the initial condition of the neuronal activity (see our full simulation details). (**h**) Box plot of the slow-variant errors across 10 training sessions for different numbers of initial conditions. (Centre lines indicate median errors, boxes indicate 25th to 75th percentiles, whiskers indicate ± 1.5× the interquartile range, and dots indicate training sessions whose error values lie outside whiskers.) (**i**) Mean error over 10 training sessions for *m* = 1, …, 10 initial conditions. (**j**) For the case of 6 initial conditions in panel (h), we plot 4 example outputs that correspond to the 5th-smallest error for the 10 training sessions. (For each simulation, we train a 400-neuron network using 40 random modulatory groups (see Methods Section 1.10).)

**Supplementary Figure 6:**
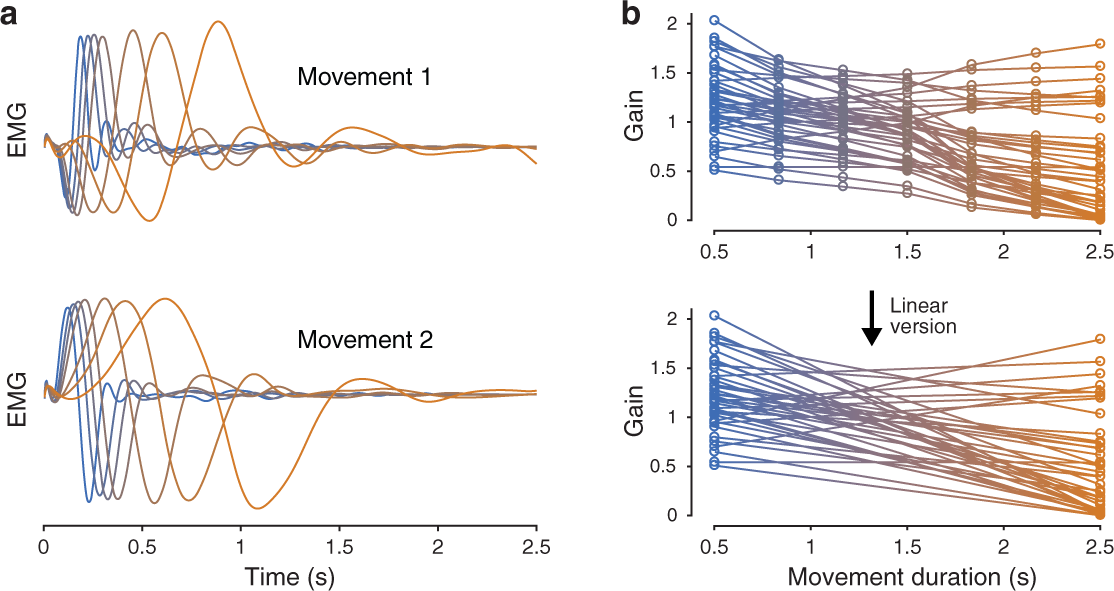
Additional results for smooth control of movement speeds through gain modulation. (**a**) We show outputs that result from the 7 trained gain patterns from Fig. 6e (which we also reproduce here in panel (b)) for both initial conditions (see Methods Section 1.10). (**b**) (Top) We reproduce the 7 optimized gain patterns for all 40 modulatory groups when training at 7 evenly-spaced speeds from Fig. 6e. We call this the ‘speed manifold’ in the main text. (Bottom) We linearly interpolate between the fast and the slow gain patterns. We use this interpolation for the outputs that we show in Fig. 6f.

**Supplementary Figure 7:**
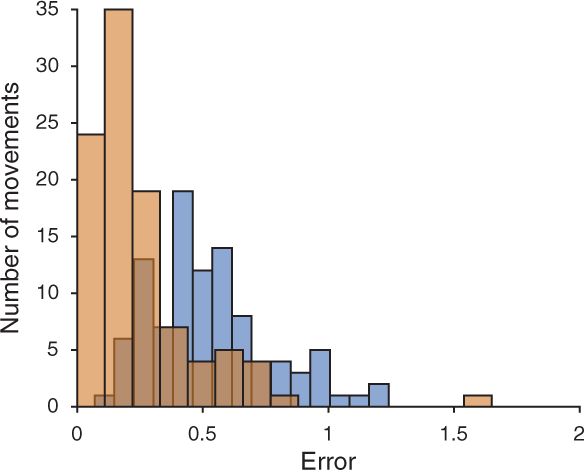
Additional results for learning gain-pattern primitives to control movement shape and speed. We plot histograms of the errors over the 100 target movements at both fast (blue) and slow (orange) speeds (see Methods Section 1.10).

**Supplementary Figure 8:**
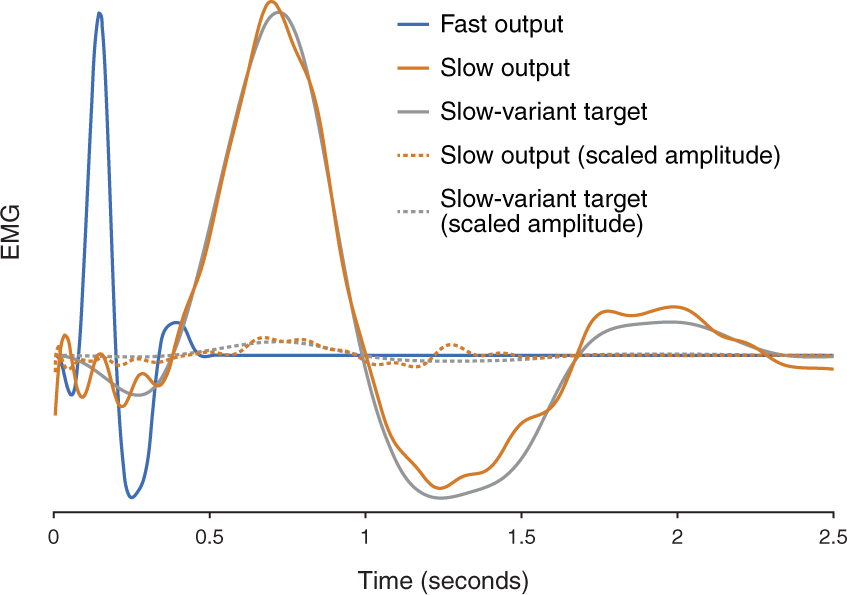
Learning slow-movement variants when scaling both the amplitude and duration of target movements. We can perform the same task as the one that we showed in Fig. 6b when we also scale the amplitude of the slow-variant target movement by the factor 1/25 (see the dashed curves). Scaling the slow-variant target movement by this factor corresponds to the same actual movement but lasting 5 times longer (see our full simulation details). We also reproduce the results from the top panel of Fig. 6b (solid curves) for comparison.

## FULL SIMULATION DETAILS

### Simulation details for Fig. 1 and Supplementary Fig. 1

We simulate two different electromyograms (EMGs) (see Methods Section 1.4) of muscle activities (initial reach and target reach) that each last 0.5 s (see Figs. 1a,f). We use a network of *N* = 200 neurons and sample transient neuronal firing rates that last 0.5 s following the initial condition ***x***_0_ of the neuronal activity (see Methods Section 1.1). We fit the readout weights over 100 trials, in which we add white Gaussian noise to the initial condition ***x***_0_ (with a signal-to-noise ratio of 30 dB) using least-squares regression so that the network output, with all gains set to 1, generates the initial reach (see Methods Section 1.5). We use the same readout weights throughout all training, and we use only one readout unit for each simulation.

In Fig. 1c, we plot the dynamics of three example neurons with all gains set to 1 (black) and all gains set to 2 (blue).

For each training iteration of the neuronal gains (to generate a target movement), we use the initial condition ***x***_0_ at time *t* = 0 (see Methods Section 1.1). We calculate the subsequent network output as described in Methods Section 1.5, and we update the neuronal gains according to Eqn. (8). We repeat this process for 18,000 training iterations (which corresponds to 2.5 hours of training time), which is enough training time for the error to saturate (see Fig. 1d).

We run 10 independent training sessions on the same target, and we plot these results in Figs. 1d,e. For each of the 10 trained gain patterns ***g***, we plot the change in the spectral abscissa of ***W*** × diag(***g***) (i.e., the largest real part in the spectrum of ***W*** × diag(***g***)) in Supplementary Fig. 1a. We observe an increase in the spectral abscissa after training. Although this change seems substantial, the resulting firing-rate activity does not change dramatically (see Supplementary Fig. 1b).

Additionally, we generate 100 network outputs for each of the 10 trained gain patterns using 100 different instances of white Gaussian noise added to the initial condition ***x***_0_ with a signal-to-noise ratio of *s* dB (where we consider values of *s* between 1 and 30 dB in increments of 1). We then calculate the square of the Euclidean 2-norm between each network output and the network output that we obtain when we do not add noise to the initial condition. We call these squared errors ***e***_1_. (This vector has 1,000 entries, with one entry for each network output.) We also generate 1,000 outputs with all gains set to 1 using 1,000 different instances of white Gaussian noise added to the initial condition *x*_0_ with a signal-to-noise ratio of *s* dB. (We again consider values of *s* between 1 and 30 dB in increments of 1.) We then calculate the square of the Euclidean 2-norm between each of these network outputs and the network output that we obtain with all gains set to 1 and no noise added to the initial condition. We call these squared errors ***e***_2_. For each signal-to-noise ratio *s*, we plot the mean and standard deviation of ***e***_1_ (i.e., the squared error corresponding to the trained gain patterns) in red and ***e***_2_ (i.e., the squared error corresponding to all gains set to 1) in blue in Supplementary Fig. 1d. We obtain very similar errors for both the trained and untrained (i.e., all gains set to 1) gain patterns, except for large (i.e., approximately larger than 25 dB) signal-to-noise ratios. For the outputs that we show in Fig. 1f, we add white Gaussian noise to the initial condition ***x***_0_ with a signal-to-noise ratio of 30 dB using one of the trained gain patterns and with all gains equal to 1.

To generate the correlation matrices that we show in Supplementary Fig. 1b, we calculate the Pearson correlation coefficient of the neuronal firing rates between all pairs of neurons in the recurrent neuronal network. Therefore, each entry in the matrix indicates the extent to which the neuronal firing rates are similar for a pair of neurons over the duration of the movement (i.e., 0.5 s). In Supplementary Fig. 1b, we show correlation matrices for examples in which all gains are set to 1 and for two example learned gain patterns. We use the same initial condition ***x***_0_ that we used during training.

We also study whether neuronal firing rates correlate more positively with a target movement after training than before training. To quantify the similarity between the neuronal firing rates and the target output, we calculate — for each of the 10 training sessions that we used in Fig. 1d — the Pearson correlation coefficient of the neuronal firing rates between each neuron and the target output. In Supplementary Fig. 1e, we plot the mean Pearson correlation coefficient across all neurons for the case in which all gains are set to 1 (i.e., before training) and for each of the 10 learned gain patterns (i.e., after training). There is a significant (with a p-value of *p* ≈ 0.002) change in the mean Pearson correlation coefficient before training versus after training using a paired Wilcoxon signed rank one-sided test. For the gain pattern that produces the largest change in the mean correlation coefficient (we show this with the grey line in the left panel of Supplementary Fig. 1e), we plot the distribution of changes in the correlation coefficients for all neurons in the bottom right panel of Supplementary Fig. 1e. We see that most values are larger than 0, so the neuronal firing rates become more positively correlated with the target output after learning. We also show an example of a substantial change in the neuronal firing rate of one neuron in the top right panel of Supplementary Fig. 1e.

In another computational experiment, we generate 10 different target muscle activities (see Methods Section 1.4) and, independently for each movement, we train either the neuronal gains, the recurrent synaptic weight matrix ***W***, the initial condition ***x***_0_, or a rank-1 perturbation of the recurrent synaptic weight matrix using a gradient-descent training procedure (with gradients that we obtain from back-propagation [53]). Before training, we use the 200-neuron stability-optimised network, initial condition ***x***_0_, and readout weights that we used in Fig. 1. Specifically, before any training, the network output is the black curve that we show in Fig. 1f. The cost function for the training procedure is the squared error between the network output and the target movement scaled by the total sum of squares of the target movement (i.e., Eqn. (7)). We run the gradient-descent training procedure until the difference between the cost function at successive training iterations is below 10^−5^ (i.e., until the cost saturates to a small value). When we train the recurrent synaptic weight matrix ***W***, after each weight update, we set any positive inhibitory weights to zero and we set any negative excitatory weights to zero. For the rank-1 perturbation, we independently train vectors ***u, v*** ∈ ℝ^200×1^ to reduce the error between the network output, which we obtain from the neuronal firing rates in Eqn. (1) with ***W*** replaced by ***W*** + ***uv***^┬^, and the target movement. Before training, the elements of ***u*** and ***v*** are chosen independently and identically from a Gaussian distribution with a mean 0 and standard deviation 0.05. In Supplementary Fig. 1f, we plot the errors for 10 different target movements for each of our 4 different training approaches.

### Simulation details for Fig. 2

For this figure, we train neuronal gains on the same task as the one that we showed in Fig. 1d — that is, we independently train 10 gain patterns to generate the target output that we showed in orange in Fig. 1f — using 3 alternative models. We use neuron-specific modulation for these simulations, as we did in Fig. 1f. We fit the readout weights so that, prior to any training (i.e., with all gains set to 1), the network output is the same in each model. (See the black curve in Fig. 1f.) We show the mean error during training in Fig. 2a. The red curve is the same error curve that we plotted in Fig. 1d, but we now use a logarithmic vertical-axis scale.

We also train the neuronal gains on the same task as above, but now using a ramping input to the network (to simulate preparatory activity prior to movement onset [4, 11]). We use the same ramping input function as the one that was used in Ref. [11]. It is exp(*t/τ*_on_) for *t* < 0 s and exp(−*t/τ*_off_) after movement onset (*t* ≥ 0), with an onset time of *τ*_on_ = 400 ms and an offset time of *τ*_off_ = 2 ms. Gain changes that result from learning now also affect the neuronal activity at *t* = 0 (i.e., at movement onset). We again run 10 independent training sessions, and we observe results that are qualitatively similar to those we saw in Fig. 1d. (See the blue curve in Fig. 2a.)

We also train a ‘chaotic’ [34] variant of our model (see Methods Section 1.3, where we describe how we construct such a model), and we train on the same target movement that we mentioned above. We use the first 0.5 s of neuronal activity. We observe a very similar error reduction over training iterations (see the grey curve in Fig. 2a) as we saw in Fig. 1d. (Compare the grey and red curves in Fig. 2a.)

Finally, we use an alternative learning rule (see Eqns. (10) and (11)) in which learning stops automatically when the difference between network outputs over successive training iterations becomes sufficiently small (see Methods Section 1.7). In Fig. 2a, we plot the error reduction using this alternative learning rule in purple. Using this alternative learning rule, we obtain a smaller error for this task (compare the purple and red curves in Fig. 2a), and learning stops after approximately 10, 000 training iterations on average.

In Fig. 2b, we plot the firing rates of 4 example neurons for each of these 4 models both before and after training the neuronal gains.

### Simulation details for Fig. 3 and Supplementary Fig. 2

For the same task as in Fig. 1, we plot the results of using random and specialized groupings (see Methods Section 1.9), as well as the neuron-specific result from Fig. 1d, in Supplementary Fig. 2a. We use the same readout weights that we used in Fig. 1.

We now give details for Figs. 3b,c and Supplementary Figs. 2b–d. We generate 5 different target outputs and run 10 independent training sessions for each target. For the random groupings (see Methods Section 1.9), we use different independently-generated random groups for each simulation. For the specialized groups (see Methods Section 1.9), for a given number of groups, we use the same grouping in all simulations. We plot the results of using 10 or 20 groups with either random or specialized groups in Figs. 3b,c and Supplementary Figs. 2b,c.

We now explain how we determine specialized groups that are shared by multiple movements (i.e., we use the same grouping for learning multiple movements); see the plots in Fig. 3c and Supplementary Figs. 2b–d. We apply *k*-means clustering (where *k* is the desired number of groups) across all of the gain patterns that we obtain using neuron-specific modulation for each of the movements. That is, we apply *k*-means clustering to a matrix of size *N* × 10 · *q*, where *N* is the number of neurons and *q* is the number of movements (and, equivalently, the number of gain patterns). We also use the specialized grouping that we obtain for 20 groups that is shared across 5 movements (see Supplementary Figs. 2b) to learn 10 hitherto-untrained movements. We plot these results in Supplementary Fig. 2d.

For the task that we just described above (i.e., training independently on 5 different target movements), we consider various different numbers of groups (using random groupings) for networks with *N* = 100,*N* = 200, and *N* = 400 neurons. We again perform 10 independent training sessions for each network, target, and number of groups. We fit the readout weights so that each scenario generates the same network output when all gains are set to 1. The readout weights remain fixed throughout training. We plot these results in Fig. 3d and Supplementary Figs. 2e–h.

We now give details for Figs. 3e,f. When we use multiple readout units, we generate 10 different initial and target outputs for each readout unit. For example, for 2 readout units, we generate 10 different initial and target outputs for each of units 1 and 2. We run independent training sessions for these 10 sets of target outputs and calculate mean errors across the 10 training sessions. For a given number of readout units, we use the same sets of initial and target outputs for all 3 network sizes and each number of random modulatory groups. We thus fit readout weights so that each scenario generates the same output with all gains set to 1. The readout weights remain fixed throughout training. We use 60,000 (instead of 18,000) training iterations to ensure error saturation.

### Simulation details for Figs. 4, 5f, and Supplementary Figs. 3,4

To create libraries of learned movements, we train a network of 400 neurons and 40 random groups (see Methods Section 1.9) on each of 100 different target movements independently. (In other words, this generates 100 different gain patterns, with one for each movement.) In Supplementary Fig. 3a, we plot the distribution of gains that we obtain after training across all 100 gain patterns. We plot all 100 outputs from these 100 learned gain patterns in Supplementary Fig. 3b. We also generate 100 new gain patterns by sampling uniformly at random from the distribution in Supplementary Fig. 3a and plot the output of each of these gain patterns in Supplementary Fig. 3c. These outputs are much more homogeneous than the learned gain patterns in Supplementary Fig. 3b, and they likely would not constitute a good basis set for movement generation.

For library sizes of *l* ∈ {1, 2, …,50}, we choose 100 samples of *l* movements (from the learned gain patterns and their outputs) uniformly at random without replacement for each *l*. We then fit the set of movements in each of the 100 sample libraries using least-squares regression for each of 100 hitherto-untrained novel target movements. We constrain the fitting coefficients *c_j_* from the least-squares regression by requiring that *c_j_* ≥ 0 for all *j* and 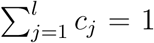. That is, we consider convex combinations of the coefficients *c_j_*. We calculate the fit error (i.e., the error between the fit and the target), the output error (i.e., the error between the output and the target), and the error between the fit and the output for each of the 100 novel movements, each of the 100 library samples, and each *l*.

For each *l* and for each randomly-generated combination of library elements (see the paragraph immediately above), we order the 100 novel target movements based on the error between the output and the fit, and we select the one that is the 50th smallest (i.e., close to the median error). We then extract the output and fit errors for this target and repeat this procedure for each of the 100 randomly-generated combinations of library elements and for *l* = 1, …,50. We plot these results in Fig. 4c and Supplementary Fig. 3g. In Fig. 4, we plot results for *l* ∈ {1, 2, …,20}; in Supplementary Fig. 3, we plot results for *l* ∈ {1,2, …,50}. Observe that there is only a small change in the errors between *l* = 20 and *l* = 50.

In Fig. 4b, for an example target (and for *l* = 2, *l* = 4,*l* = 8, and *l* = 16), we plot the output and fit that produce the 50th-smallest error between the output and the target across the 100 randomly-generated libraries. In Supplementary Fig. 3e, we calculate the median error over the 100 target movements and we plot the distribution of these median errors over the 100 randomly-generated combinations of library elements for *l* = 5 and *l* = 20.

Additionally, for each *l* and for each of the 100 target movements, we order the 100 combinations of library elements based on the error between the output and the fit, and we select the one that is the 50th smallest. We then extract the output and fit errors for this combination and repeat this procedure for each of the 100 target movements and for *l* = 1, …,50. We plot these results in Supplementary Fig. 3h. This indicates that we obtain qualitatively similar results if we average over the 100 target movements or if we instead average over the 100 combinations of library elements. In Fig. 4d and Supplementary Fig. 3d, we first calculate the median error over the 100 target movements for each *l* and for each of the 100 combinations of library elements. We then plot the median of these errors over the 100 combinations of library elements for each *l*.

We also calculate the Pearson correlation coefficient between the output and the fit errors for each *l* when taking the 50th-smallest error across the 100 novel target movements (see Supplementary Fig. 3i) or across the 100 randomly-generated samples (see Supplementary Fig. 3j).

We also repeat these simulations for the baseline rate *r*_0_ = 5 Hz in Eqn. (2). We plot the results of these simulations in Fig. 5f and Supplementary Fig. 4, and we note that we obtain very similar results to those that we obtained for *r*_0_ = 20 Hz.

### Simulation details for Figs. 5a–e

We now describe the details of our simulations when using a baseline rate of *r*_0_ = 5 Hz.

For the 200-neuron network that we used in Fig. 1, we plot the (relative to baseline) firing rate *f*(*x_i_*) (see Eqn. (2)) of 20 excitatory and 20 inhibitory neurons in Fig. 5 with (panel (a)) *r*_0_ = 20 Hz in Eqn. (2) and (panel (b)) *r*_0_ = 5 Hz. In Fig. 5c, we plot the relative firing rate of all neurons over time versus the relative firing rate when using the linear gain function *f*(*x_i_; g_i_*) = *g_i_x_i_* for the cases of (black) *r*_0_ = 20 Hz and (blue) *r*_0_ = 5 Hz. We set all of the gains to 1 for these simulations.

We also train a recurrent neuronal network on the same task as the one that we showed in Figs. 1d–f, except with a baseline rate of *r*_0_ = 5 Hz. We plot these results in Figs. 5d–e and compare them to our observations for *r*_0_ = 20 Hz. For the 10 noisy initial conditions that we used to generate the outputs in the inset in Fig. 5d, we add white Gaussian noise to the initial condition *x*_0_ with a signal-to-noise ratio of 30 dB. In other words, we generate noise in the same manner as we did in Fig. 1f.

### Simulation details for Fig. 6 and Supplementary Figs. 5,6,8

We now describe our simulations for learning target activity that lasts longer than 0.5 s. In each of these simulations, we use a network of 400 neurons and 40 random modulatory groups. (See Methods Section 1.9 for details on how we determine such groups.) We construct ‘slow’ (2.5 s) target movements with *σ* = 550 ms and *ℓ* = 250 ms in Eqn. (5). We then construct a ‘fast’ (0.5 s) variant of each movement. Each movement variant has 500 evenly-spaced points (see Methods Section 1.4). We sample the fast variant using 100 evenly-spaced points, and we then augment 400 instances of 0 values to the final 2 s of the movement to ensure that both movement variants have the same length. (See the top right of Fig. 6a.)

#### Details for Fig. 6b, Supplementary Figs. 5a,c,e, and Supplementary Fig. 8

For Fig. 6b, we fit readout weights using least-squares regression, such that with all gains set to 1, the network output approximates the fast variant. We then train gain patterns using our learning rule in Eqns. (8) and (9) so that the network output generates the slow-movement variant. (The initial condition *x*_0_ and readout weights remain fixed.) We use 60,000 training iterations, and we run 10 independent training sessions for each of 10 different target movements. We plot one such movement in Fig. 6b, and we plot results of all simulations in Supplementary Figs. 5a,c. For Supplementary Fig. 8, we perform the same task except that we scale the amplitude of the slow-movement variant by the factor 1/25. Scaling the slow-variant target movement by this factor corresponds to the same actual movement but lasting 5 times longer (see above). In Supplementary Fig. 8, we show results for the same example that we plotted in Fig. 6b.

#### Details for Fig. 6c and Supplementary Figs. 5b,d,f,g

We wish to obtain neuronal dynamics that are less sensitive to noisy initial conditions than those that we generated from gain patterns that we obtained from our learning rule (i.e., those that we plot in Supplementary Figs. 5a). For example, in Fig. 6b, the neuronal firing rates have decayed substantially towards baseline after approximately 0.75 s, even though the output activity is close to its maximum value. Therefore, a small change in the initial condition would likely substantially affect the neuronal activity for times after approximately 0.75 s. We therefore perform the task that we described in the paragraph above (i.e., generating a slow-movement variant by changing neuronal gains) using a gradient-descent training procedure with gradients that we obtain from back-propagation [53]. Together with learning the gain pattern for the slow variant, we jointly optimize a single set of readout weights (shared by both the fast-movement and slow-movement variants), as we discussed in Methods Section 1.5, as part of the same training procedure. The gains are still fixed at 1 for the fast variant. The cost function for the training procedure is equal to the squared Euclidean 2-norm between actual network outputs and the corresponding target outputs at both fast and slow speeds plus the Euclidean 2-norm of the readout weights, where the latter acts as a regularizer. We run gradient descent for 500 iterations, which is well after the cost has stopped decreasing.

Using the target movement from Fig. 6b, we plot the output of the back-propagation training procedure in Fig. 6c, and we plot results of all simulations in Supplementary Figs. 5b,d on the same 10 target movements as those that we used in Supplementary Fig. 5a. In Supplementary Fig. 5g, for the outputs in Figs. 6b,c, we add white Gaussian noise with a signal-to-noise ratio of 4 dB to the initial condition. We observe that the outputs from the back-propagation training procedure are less sensitive than the outputs from the learning rule to noisy initial conditions.

#### Details for Supplementary Figs. 5h–j

In these simulations, we train a single gain pattern that is shared by *m* different movements, which each last 2.5 s and where each movement corresponds to a different initial condition (IC). To generate a collection of *m* such ICs, in which each IC evokes neuronal activity of approximately equal amplitude with all gains set to 1, we randomly rotate the top *m* eigenvectors of the observability Gramian of the matrix ***W*** − ***I*** [11]. Specifically, we do this by creating a matrix of *m* columns — one for each of these *m* eigenvectors — and right-multiplying this matrix by a random *m × m* orthogonal matrix (which we obtain via a QR decomposition of a random matrix with elements drawn from a normal distribution with mean 1 and standard deviation 1).

Given *m* ICs, we uniformly-at-random choose *m* fast target movements and their slow counterparts out of a fixed set of 10 different movements. We then train a recurrent neuronal network to generate the correct fast and slow target movements by optimizing a single set of readout weights (shared by both fast and slow variants) and a single gain pattern that generates the slow variants (where we set the gains for each of the fast variants to 1). We train using the same gradient-descent method with back-propagation that we described above for Fig. 6c. We plot the results as a function of the number *m* of movement–IC pairs (see Supplementary Figs. 5h,i) for 10 independent draws of the ICs that we just described above.

#### Details for Fig. 6d; top panel

For each of the 10 trained movements in Supplementary Figs. 5a,b, we extract the mean minimum error across all simulations for the outputs that we obtain both from our learning rule (see Supplementary Fig. 5a) and from training via back-propagation (see Supplementary Fig. 5b). We then linearly interpolate between the learned gain patterns for the fast and slow outputs, and we and calculate the error (see Methods Section 1.6) between the output and the target movement at the interpolated speed. We calculate these errors for many interpolated movement durations between 0.5 s and 2.5 s, and we plot the mean errors for both our learning rule and the back-propagation training in the top panel of Fig. 6d. We also show an example output that lasts 1.5 s.

#### Details for Figs. 6d–f and Supplementary Fig. 6

To demonstrate that gain modulation can provide effective smooth control of movement speed for multiple initial conditions of the neuronal activity, we train networks to generate a pair of target movements in response to a corresponding pair of orthogonal initial conditions (see the above description of Supplementary Figs. 5h–j) at fast and slow speeds and also at each of 5 intermediate, evenly-spaced speeds in between these extremes. To do this, we parametrize the gain pattern of speed index *s* (with *s* ∈ {1, …,7}) as a convex combination of a gain pattern ***g***_*s*=1_ for fast movements and a gain pattern ***g***_*s*=7_ for slow movements, with interpolation coefficients of *λ_s_* (with ***g***_*s*_ = *λ*_*s*_***g***_*s*=1_ + (1 − *λ_s_*)***g***_*s*=7_, *λ*_1_ = 1, and *λ*_7_ = 0). We optimize (using back-propagation, as discussed above) over ***g***_*s*=1_, ***g***_*s*=7_, the 5 interpolation coefficients *λ_s_* (with *s* ∈ {2, …,6}), and a single set of readout weights. For a given speed *s*, we use the gain pattern ***g***_*s*_ for both movements.

We plot the 7 learned gain patterns in Fig. 6e, and we plot their corresponding outputs for both initial conditions in Supplementary Fig. 6. (We call this collection of 7 trained gain patterns the ‘speed manifold’.) We show the linear version of the speed manifold (i.e., interpolating between the fast and slow gain patterns) in Supplementary Fig. 6b. Interpolating between the fast and slow gain patterns accurately generates both movements at any intermediate speed. (See the bottom panel of Fig. 6d.). For both initial conditions, we plot outputs at 5 evenly-spaced speeds by linearly interpolating between the fast (***g***_*s*=1_) and slow (***g***_*s*=7_) gain patterns in Fig. 6f.

### Simulation details for Fig. 7

We simultaneously train gain patterns for controlling different movements (i.e., different movement shapes) and their speed. We train a recurrent neuronal network (using back-propagation, as we discussed previously) to generate each of 10 different movement shapes at 7 different, evenly-spaced speeds (ranging from the fast variant to the slow variant) using a single fixed initial condition *x*_0_. To jointly learn gain patterns that control movement shape and speed, we parametrize each gain pattern as the element-wise product of a gain pattern that encodes shape (which we use at each speed for a given shape) and a gain pattern that encodes speed (which we use at each shape for a given speed). We again parametrize (see our details for Figs. 6d–f) the gain pattern that encodes the speed index *s* (with *s* ∈ {1, …,7}) as a convex combination of two common endpoints, ***g***_*s*=1_(which we use for the fast-movement variants) and ***g***_*s*=7_ (which we use for the slow-movement variants). We thus optimize over 10 gain patterns for movement shape, 2 gain patterns each for fast and slow movement speeds, 5 speed-interpolation coefficients (see above), and a single set of readout weights.

In Fig. 7b, we plot the gain patterns that we obtain for controlling the movement speeds at each of the 7 trained speeds. In Fig. 7c, we show the mean error between the network output and the target over the 10 target movements when generating gain patterns for movement speed by linearly interpolating between the trained fast (***g***_*s*=1_) and slow (***g***_*s*=7_) gain patterns. In Fig. 7d, we plot the outputs of 6 of the 10 gain patterns for movement shape at each of 5 interpolated speeds between the fast and the slow gain patterns. In rightmost panel of Fig. 7a, we plot 2 example movement shapes at 3 interpolated speeds.

### Simulation details for Fig. 8 and Supplementary Fig. 7

For these figures, we use the 10 trained gain patterns for movement shapes, as well as the speed manifold from Fig. 7 (see our simulation details for Fig. 7). Using our learning rule from Eqns. (8) and (9), we train 10 coefficients *c*_1_, …, *c*_10_ (with one for each shape-specific gain pattern; see Fig. 8a) to construct a new gain pattern that, together with the speed manifold, generates a new target movement at the fast and slow speeds. Specifically, we replace the gains *g*_*i*_ (for *i* ∈ {1, …, *N*}) with the coefficients *c_i_* (for *i* ∈ {1, …,10}) in Eqns. (8) and (9). We use the mean of the errors at the fast and slow speeds. To generate the network output at the fast and slow speeds, respectively, we calculate the element-wise product between the newly-constructed gain pattern and the fast and slow gain pattern, respectively, on the speed manifold. We independently train, using 10,000 training iterations, the coefficients *c*_1_, …, *c*_10_ on each of the 100 target movements that we used for Fig. 4. In Supplementary Fig. 7, we plot histograms of the errors over the 100 target movements after training for both the fast and slow speeds. We plot the mean error (see the black curve) over all 100 target movements at interpolated speeds in Fig. 8c. For the output that produces the 50th-smallest summed errors from fast and slow speeds, we plot the error in red in Fig. 8c. As a control, we calculate the mean error between the network output and the target over the 100 target movements when choosing one of the 100 newly-learned gain patterns uniformly at random without replacement. (See the grey curve in Fig. 8c.)

Additionally, instead of learning to combine gain patterns using the method that we described in the previous paragraph, we determine coefficients *c*_1_, …, *c*_10_ using a least-squares regression by fitting the 10 learned movements to each of the 100 target movements at the fast and slow speeds simultaneously and requiring that *c_j_* ≥ 0 for all *j* and 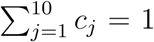. (See the black dashed curve in Fig. 8c.)

Finally, we plot the Pearson correlation coefficient between pairs of target movements versus the Pearson correlation coefficient between corresponding pairs of learned coefficients *c*_1_, …, *c*_10_. In our visualization, we plot only 1,000 of the 4,950 data points. (We choose these points uniformly at random.) Note that we are unlikely to observe correlation values close to −1 between pairs of combination coefficients because the coefficients *c*_1_, …, *c*_10_ are likely to form a convex combination (see our discussion of Fig. 4); in fact, we calculate the mean sum of the coefficients to be approximately 0.91.

